# An image-computable model for the stimulus selectivity of gamma oscillations

**DOI:** 10.1101/583567

**Authors:** Dora Hermes, Natalia Petridou, Kendrick Kay, Jonathan Winawer

## Abstract

Gamma oscillations in visual cortex have been hypothesized to be critical for perception, cognition, and information transfer. However, observations of these oscillations in visual cortex vary widely; some studies report little to no stimulus-induced narrowband gamma oscillations, others report oscillations for only some stimuli, and yet others report large oscillations for most stimuli. To reconcile these findings and better understand this signal, we developed a model that predicts gamma responses for arbitrary images and validated this model on electrocorticography (ECoG) data from human visual cortex. The model computes variance across the outputs of spatially pooled orientation channels, and accurately predicts gamma amplitude across 86 images. Gamma responses were large for a small subset of stimuli, differing dramatically from fMRI and ECoG broadband (non-oscillatory) responses. We suggest that gamma oscillations in visual cortex serve as a biomarker of gain control rather than being a fundamental mechanism for communicating visual information.

## Introduction

An important goal in visual neuroscience is to develop models that can predict neuronal responses to a wide range of stimuli. Such models are a test of our understanding of how the system functions, and have led to insights about canonical computations performed by the early visual system, such as filtering, rectification, and normalization (Carandini et al., 2005). Image-computable models, which predict responses to arbitrary, unlabeled images, have been developed for the functional MRI (fMRI) blood oxygen level dependent (BOLD) signal (Dumoulin and Wandell, 2008; Güçlü and van Gerven, 2015; Kay et al., 2013a, 2013b) and for spiking of single neurons in animals (Mante et al., 2008; Rust et al., 2005; Simoncelli and Heeger, 1998). In contrast, to our knowledge, there are no image-computable models to predict oscillations in the gamma band (30-80 Hz) of the local field potential, although gamma oscillations have been intensely studied and are a prominent part of many neural recordings and theories of neural function (e.g. (Jensen et al., 2007; Tallon-Baudry and Bertrand, 1999)).

Studies in humans and animal models indicate that gamma oscillations are systematically related to image properties, supporting the possibility that this stimulus dependence might be captured in an image-computable model. In particular, oriented bars (Gray and Singer, 1989; Gray et al., 1989) and gratings (Eckhorn et al., 1988; Hermes et al., 2017a; Muthukumaraswamy and Singh, 2008a) elicit large gamma responses in visual cortex. The responses tend to increase with stimulus size and contrast (Gieselmann and Thiele, 2008; Henrie and Shapley, 2005; Ray and Maunsell, 2011), and tend to decrease in the presence of unstructured noise (Bartoli et al., 2019; Hermes et al., 2015; Jia et al., 2013; Kayser et al., 2003) or multiple orientations (Bartolo et al., 2011; Lima et al., 2010). Some images of scenes and objects cause large reliable gamma oscillations while others do not (Brunet et al., 2015; Hermes et al., 2015). Gamma response selectivity differs from that of the BOLD response (Hermes et al., 2017b; Muthukumaraswamy and Singh, 2008b) and single-and multi-unit spike rates (Jia et al., 2013; Peter et al.; Ray and Maunsell, 2011). For example, when recording from the same electrode in V1, increasing grating size causes gamma oscillations to increase in power while causing firing rates to decrease (Jia et al., 2013; Ray and Maunsell, 2011). Gamma power also decreases dramatically with noise masking while firing rates do not change substantially (Jia et al., 2013). Because image selectivity in gamma oscillations clearly differs from the selectivity in firing rates and BOLD signals, a model to predict the extent to which different images will give rise to gamma oscillations requires a different form than a model to predict the BOLD signal or firing rates.

Here, we measured responses from electrocorticography (ECoG) electrodes over visual cortex while human patients viewed a variety of different images. We separated the ECoG response into two spectrally overlapping components: one broadband (spanning 30-200 Hz) and one narrowband (centered between 30-80 Hz). We compared the broadband component and narrowband gamma component to the images, and developed image-computable models to account for the stimulus selectivity present in each. The broadband response was well fit by a model adapted from fMRI of visual cortex (Kay et al., 2013a). The narrowband gamma responses were strikingly different, and we developed a new, image-computable model in order to explain those responses. The differences in the patterns of responses and the differences in the two models suggest that broadband signals and narrowband gamma originate from distinct aspects of neural circuitry.

## Results

Narrowband gamma power is highly stimulus dependent. For example, there are large gamma oscillations in electroencephalography (EEG) (Murty et al., 2018; Scheeringa et al., 2011), magnetoencephalography (MEG) (Hoogenboom et al., 2006; Muthukumaraswamy and Singh, 2008b, 2008a), and microelectrode local field potentials (LFP) (Gray and Singer, 1989; Gray et al., 1989) responses to bars and gratings, but not to noise patterns (Bartoli et al., 2019; Hermes et al., 2015; Jia et al., 2013) nor many natural images (Hermes et al., 2015; Kayser et al., 2003). To our knowledge, there is no model that can predict the amplitude of gamma oscillations elicited by arbitrary, unlabeled images. Here, we recorded ECoG data from three subjects viewing 86 static, band-passed, grayscale images and developed an image-computable model to predict the level of gamma oscillations observed for these 86 stimuli.

We identified ECoG electrodes that were located on the surface of V1, V2 and V3 and had a well-defined population receptive field (pRF) measured from an independent experiment with sweeping bar stimuli (as in (Winawer and Parvizi, 2016; Winawer et al., 2013)). This yielded 6 electrodes in the first subject, 2 in the second subject and 7 in the third subject (Figure 1A). For each electrode we calculated the power spectra from the 500-ms window following presentation of each of the 86 images. As in previous work (Hermes et al., 2015), we separated the power spectra into an oscillatory and non-oscillatory component by modeling the log-power/log-frequency spectrum as the sum of 3 components: a linear baseline, a constant, and a Gaussian centered between 30 and 80 Hz (Figure 1B). These three terms correspond to the baseline signal in the absence of a stimulus, a stimulus-specific broadband response, and a stimulus-specific narrowband gamma response, respectively. Note that our quantification of narrowband gamma response was constrained to be nonnegative, and so in the presence of noise, even images which cause the oscillatory gamma response to decrease or to remain unchanged (compared to the blank screen) are likely to be estimated as slightly positive. Replicating previous results (Hermes et al., 2015; Jia et al., 2013; Zhou et al., 2008), we observe large increases in narrowband gamma power for large, high-contrast grating patterns but not for noise patterns, and strong broadband responses for both grating patterns and noise stimuli (Figure 1B-C). Because the broadband response is similar for the two types of stimuli, the lack of narrowband gamma for the noise stimuli does not indicate a general lack of visual responsivity, but rather that a different type of stimulus selectivity exists for the gamma response compared to the broadband response.

**Figure 1:**
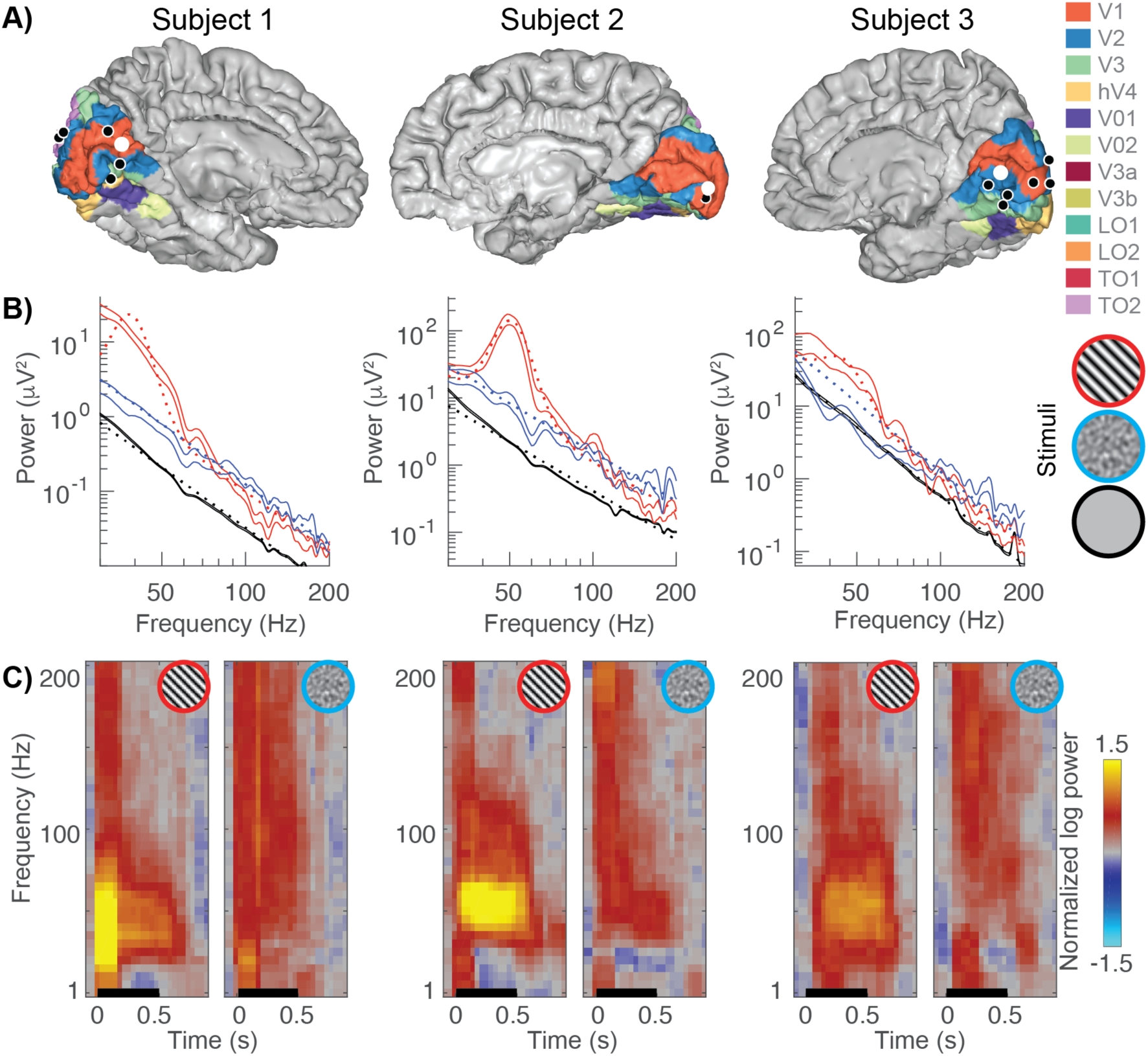
Example broadband and gamma responses to grating and noise stimuli. **A)** Location of the electrodes implanted in each subject (black and white dots) rendered on estimates of early visual areas (Benson and Winawer, 2018; Benson et al., 2012). **B)** The power spectra for example electrodes (white dots from panel A). Power spectra are shown on a double logarithmic plot for a grating stimulus (red, stimulus number 45), a noise pattern (blue, stimulus number 83) and the baseline condition (black). The solid lines indicate the data (68% confidence interval from bootstrapping). The dotted lines indicate the fits to the data: stimulus-induced responses are modeled as a baseline linear fit (black) plus a constant and Gaussian to capture broadband and narrowband stimulus-specific responses, respectively. **C)** Time-frequency plots (spectrograms) for the same electrodes. The black line indicates stimulus timing (500 ms). All spectrograms are normalized with respect to the same baseline: the inter-stimulus interval between all trials (from 250–500 ms after stimulus offset). Spectrograms are cut off at a maximum power of ±1.5 log10 units. The multitaper approach results in a temporal smoothing of 200 ms and a frequency smoothing of ±15 Hz. Spectrograms represent averages across all trials of a given stimulus type.

### Stimulus selectivity differs for ECoG gamma oscillations compared to broadband ECoG and BOLD

We find that stimulus selectivity is similar for broadband ECoG and BOLD but quite different for gamma oscillations. We first illustrate this with example responses from the 3 signal types. BOLD responses to a subset of these images were measured in previous work (Kay et al., 2013a) (images 1 to 77, Figure 2A). Using the publicly available data from that study (http://kendrickkay.net/socmodel/), we identified a voxel in V1 with a similar pRF location to one of our electrode’s pRF (electrode 3 in subject 1), and plotted the BOLD responses from this voxel to the different stimuli (Figure 2B). The pattern of responses in the BOLD data is generally similar to that of the broadband ECoG data (Figure 2C). In contrast, the ECoG narrowband gamma responses (Figure 2D) are quite different. The most salient pattern across the 3 types of responses to the whole image set is substantial BOLD and broadband responses for most images whereas gamma responses are large only for gratings (*Δorientation*). The BOLD signal and ECoG broadband power are largest for patterns with multiple orientations, such as the curvy patterns in images 74-78 (and similar patterns from the *Δspace* stimuli that overlapped the pRFs, including images 8-16 and 30-36). These responses often exceeded the response to high contrast gratings (images 39-46). The opposite is true for the gamma responses, for which the biggest response by far is to high-contrast gratings. For all three signal types, if there is a response to a stimulus of a particular pattern, such as gratings (images 47-50) or plaids (images 51-54), then responses increased with stimulus contrast for that pattern. Note that the scale of the different measures varies considerably (e.g., ∼5% signal change for BOLD, 200% for broadband, 2,000% for gamma) and are not comparable. We return to this issue in the Discussion.

**Figure 2.**
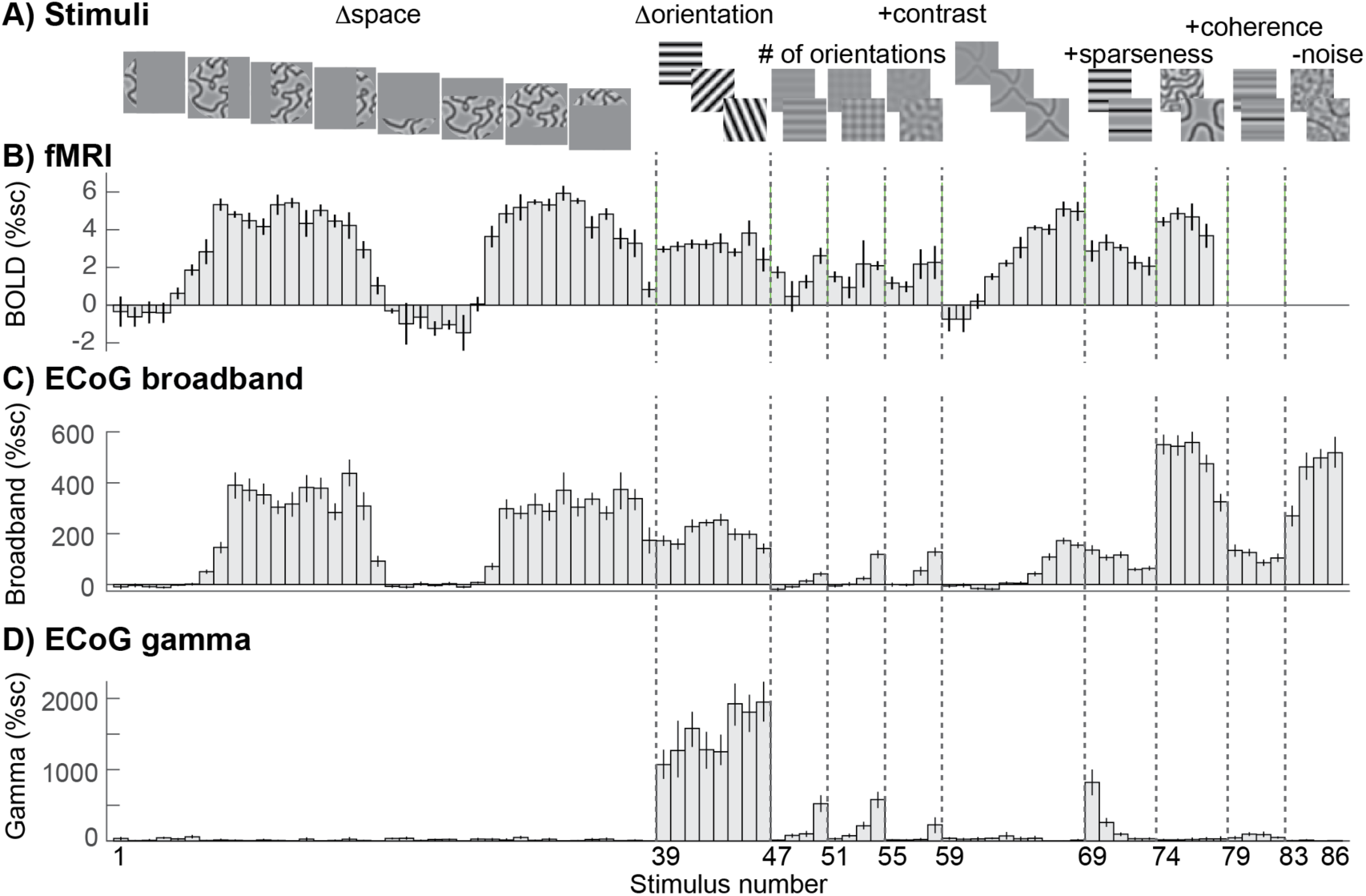
Stimulus selectivity for BOLD, broadband, and gamma. Responses in human V1 to different images. **A)** The images were identical to those used in a prior fMRI study, with a few additions, and are grouped into several stimulus categories: SPACE (1 to 38), ORIENTATION (39 to 46), CONTRAST (47 to 68), SPARSITY (69 to 78), COHERENCE (79 to 86). **B)** The fMRI BOLD response in one V1 voxel whose pRF location is matched to the electrode shown in C and D (replotted from (Kay et al., 2013a)). **C)** The ECoG broadband response for all 86 images (recorded from the same electrodes as shown in Figure 1 - left). **D)** The ECoG narrowband gamma response for all 86 images, recorded in the same electrode as C). All plots show percent signal change, and error bars display the 68% range for bootstrapped responses. All images are shown in Supplemental Figures S1-S5.

These patterns are clearly evident not just in the example electrode (Figure 2), but also in the responses averaged across the 15 electrodes in V1-V3 of 3 subjects (Figure 3). In particular the broadband responses to curved patterns (stimuli 1-39) are high whereas the gamma responses are relatively low. Another systematic difference between the two types of responses is that broadband increases with the number of component gratings, whereas narrowband gamma decreases with the number of component gratings (Figure 3 insets, showing the responses to stimuli made from 1 (grating), 2 (plaid), or 16 (circular) component gratings).

**Figure 3.**
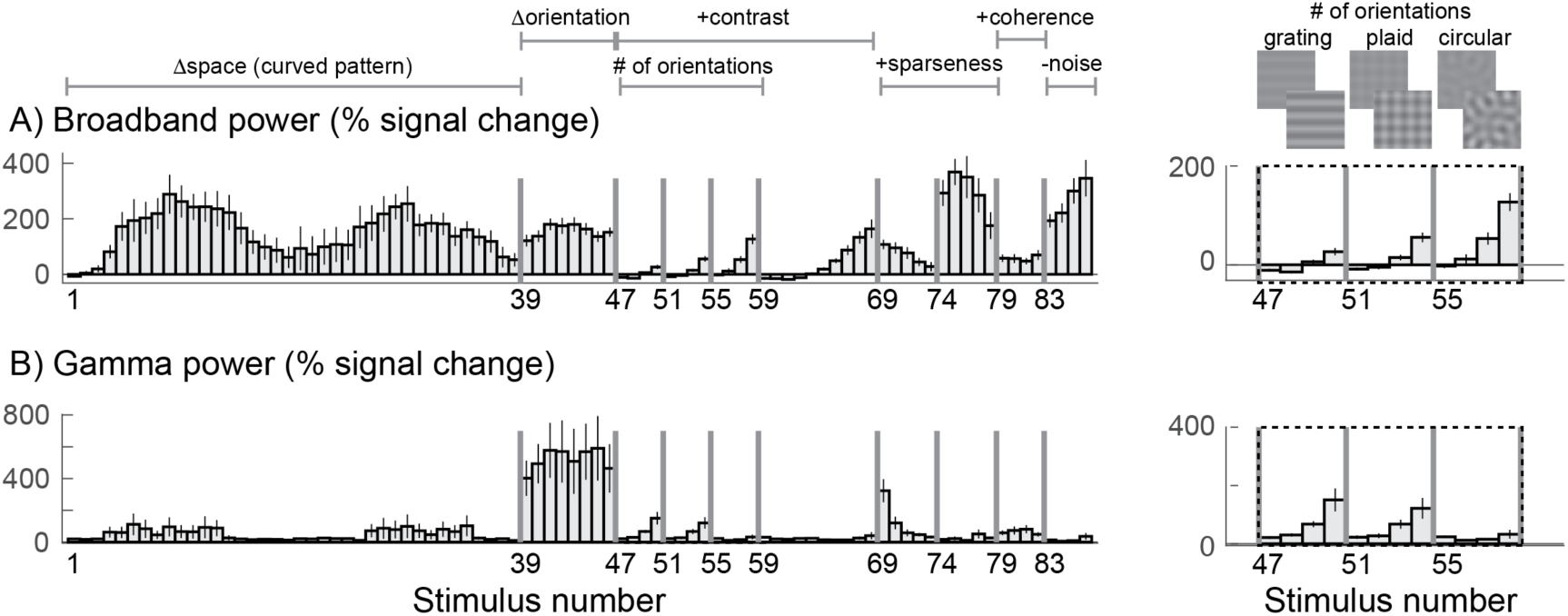
Stimulus selectivity for broadband and gamma across 15 electrodes in V1-V3. Broadband (A) and narrowband gamma (B) responses to 86 stimuli. Data are plotted as in Figure 2, except that bars represent the mean across 15 electrodes in V1-V3, and error bars represent the 68% confidence interval from bootstrapping across the electrodes. Insets (right) show a zoomed-in view of how the responses vary with 4 contrast levels and different numbers of component gratings: 1, ‘grating’, 2 ‘plaid’, or 16 ‘circular’. Within each type of pattern, the 4 bars are responses to stimuli with increasing stimulus contrast.

### Broadband changes are well predicted by a model developed for fMRI

A variety of models have been developed to predict visually evoked fMRI signals, ranging from simple linear isotropic pRF models (Dumoulin and Wandell, 2008) to high-dimensional filter models with thousands of basis functions (Eickenberg et al., 2017; Güçlü and van Gerven, 2015; Kay et al., 2008) to cascade models composed of a small number of canonical computations (Kay et al., 2013a). (For a review see (Wandell and Winawer, 2015)). Since ECoG broadband responses typically correlate well with BOLD in visual cortex (Hermes et al., 2017b; Winawer et al., 2013), we tested whether a second-order contrast model (SOC) developed for fMRI could accurately fit, and predict, the ECoG broadband signal (Kay et al., 2013a). The SOC model is a two-stage LNN (Linear-Nonlinear-Nonlinear) cascade model. The first stage includes filtering the images with oriented Gabor functions (L), rectification (N), and divisive normalization (N). The second stage includes spatial summation within a pRF (L), calculation of second-order contrast (N) and a compressive nonlinearity (N). Figure 4 shows the ECoG broadband responses for two example electrodes in each of the 3 patients measured in V1, V2 or V3 (all electrodes are shown in Supplemental Figures S6-S7). The SOC model was fit to the broadband changes, using leave-one-stimulus-out cross-validation to control for overfitting and to obtain unbiased estimates of model performance. Although there are some discrepancies (especially in stimuli 79-86), the model fit the data well overall and explained an average of 82% of the cross-validated variance across the 15 ECoG electrodes.

**Figure 4.**
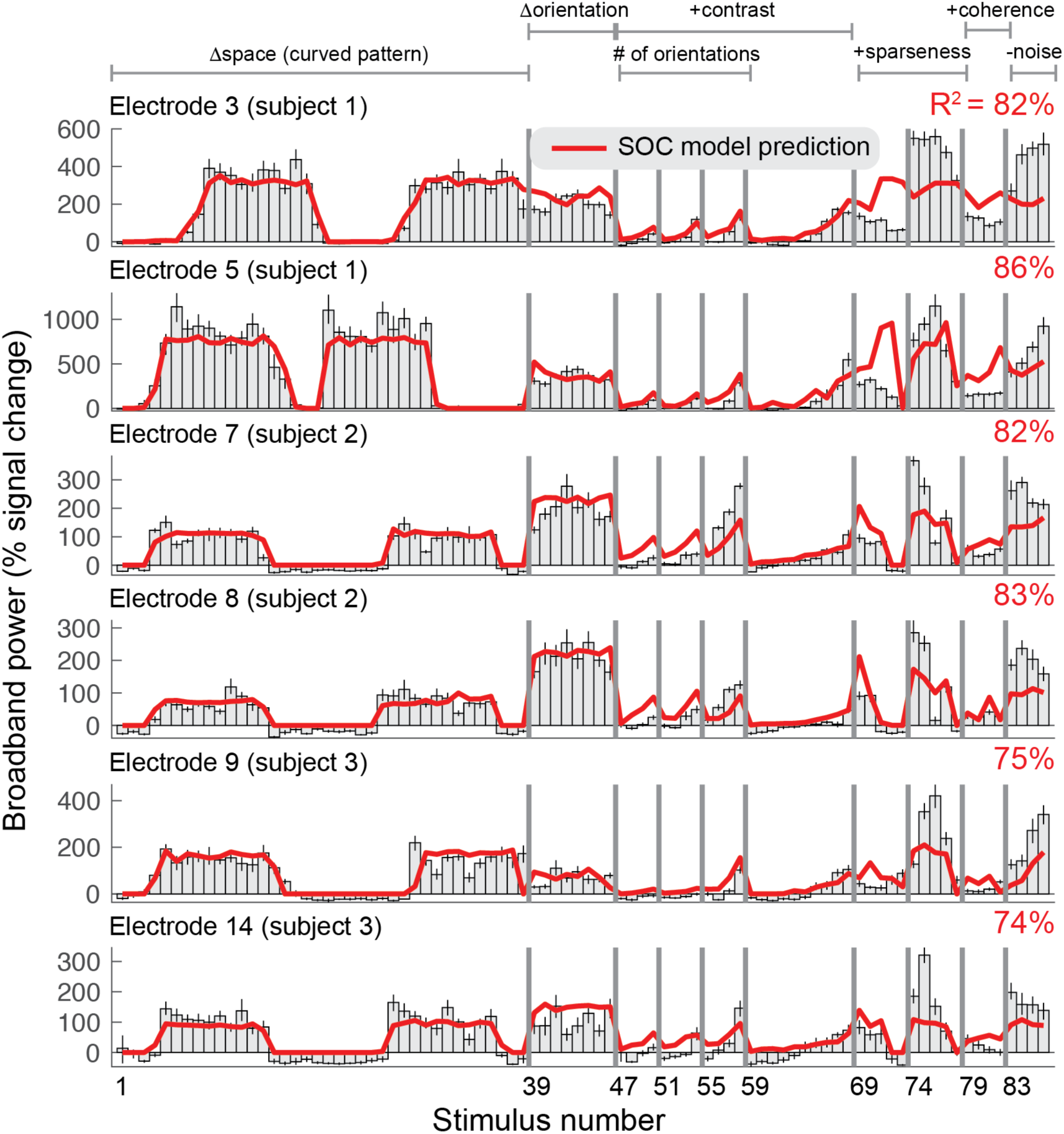
Second-order contrast (SOC) model accounts for ECoG broadband responses. Each row shows the percent signal change in ECoG broadband power for all 86 stimuli for six of the 15 electrodes on V1, V2 or V3. Error bars display the 68% range for bootstrapped responses (bootstrapped across repeated presentations of the same stimuli). The SOC model was fit to these data using leave-one-stimulus-out cross-validation. The cross-validated predictions and amount of variance explained are shown in red.

### An image-computable model of narrowband gamma responses

The striking difference in the selectivity observed in gamma responses compared to BOLD and broadband motivated us to develop a new model. Whereas the SOC model predicts increased responses for images with multiple orientations (e.g., plaids compared to gratings) and for sparse patterns compared to gratings, the gamma selectivity exhibits the opposite pattern. We propose a new model, called the Orientation Variance (OV) model, that computes variance across the outputs of spatially pooled orientation channels within a pRF. Similar to the SOC model, the OV model builds on known computations in the visual cortex, but differs in the way signals are pooled across orientation bands. In particular, the model first sums outputs across space within an orientation band. This computation can be thought of as an analog to the long-range horizontal connections in visual cortex, which preferentially connect cortical columns tuned to the same orientation (Gilbert and Wiesel, 1983; Hata et al., 1988).

In the OV model, images are first filtered with oriented Gabor patches and combined across quadrature phase (contrast energy), as in the first stage in the SOC model (Figure 5A). This results in 8 oriented contrast-energy images, one image per orientation band. However, unlike the SOC model, the OV model then sums the contrast energy of each of the orientation images separately within the pRF, resulting in 8 values, one for each orientation band (Figure 5B). The motivation for this is that gamma oscillations are sensitive to images dominated by a single orientation and are sensitive to the size of the image; pooling within a band preserves information about the response level of each band summed over space. This is in contrast to the SOC model of BOLD and broadband, which combines the local responses across orientation bands first and then pools over space. Finally, variance across these 8 values is calculated and a power-law nonlinearity (exponent *n*) and a multiplication with a gain (*g*) were included to predict the ECoG response (Figure 5C).

**Figure 5.**
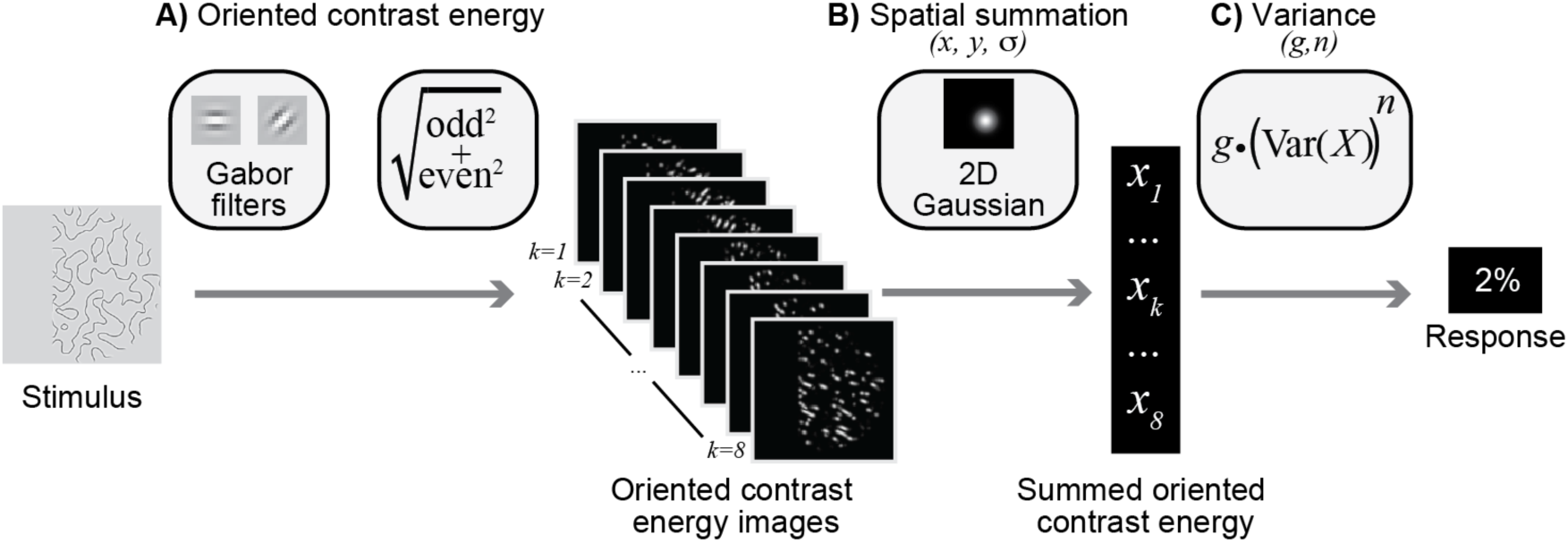
Orientation Variance (OV) model of gamma responses. In the OV model, responses are largely driven by contrast and variance across orientations in the population receptive field. **A)** Oriented contrast energy. Images are filtered with quadrature pair Gabor filters occurring at 8 orientations. The quadrature pairs are summed across phase, resulting in 8 images with contrast energy for each orientation. **B)** The contrast energy within each of these 8 images is summed within a population receptive field defined by a Gaussian with parameters *x, y* and *σ*. This results in 8 values, indicating the summed contrast energy within the pRF for each orientation. **C)** Variance is calculated across these 8 values, followed by a power-law nonlinearity (*n*) and a gain (*g*). Intuitively, the model predicts a large response when only one or a few orientations have high contrast energy and a low response when all orientations have similar contrast energy.

We illustrate the model behavior with a few simple texture patterns (for model parameters *n* = 0.5 and *g* = 1). The OV model predicts a large response when variance across orientations is high and a small response when variance is low. For simplicity, we assume the pRF is as large as the image patch. For a grating pattern, one orientation channel has a large output (high contrast energy), the two neighboring channels have medium outputs, and the 5 others have small outputs (top left Figure 6). The variance across the 8 summed contrast energies will thus be high. When the grating is reduced in contrast (bottom left Figure 6), the relative outputs of the orientation bands are unchanged, but the variance is lower. If a second, perpendicular grating is overlaid on the first to create a plaid pattern, matched to the grating in total root mean square (RMS) contrast energy (summed across bands), variance is lower (top right Figure 6), and hence the predicted signal is lower. Finally, for a pattern with many orientations at approximately equal contrast, such as in the case of the curved patterns (bottom right Figure 6), the variance will be quite low. The OV model thus has two important properties: first, the output increases with contrast within the pRF (high contrast gratings produce higher responses than low contrast gratings), and second, the output increases with increasing variance across orientated contrast energy within the pRF (gratings produce higher responses than plaids, and plaids produce higher responses than curved patterns, consistent with the observed data in Figure 3b, insets).

**Figure 6.**
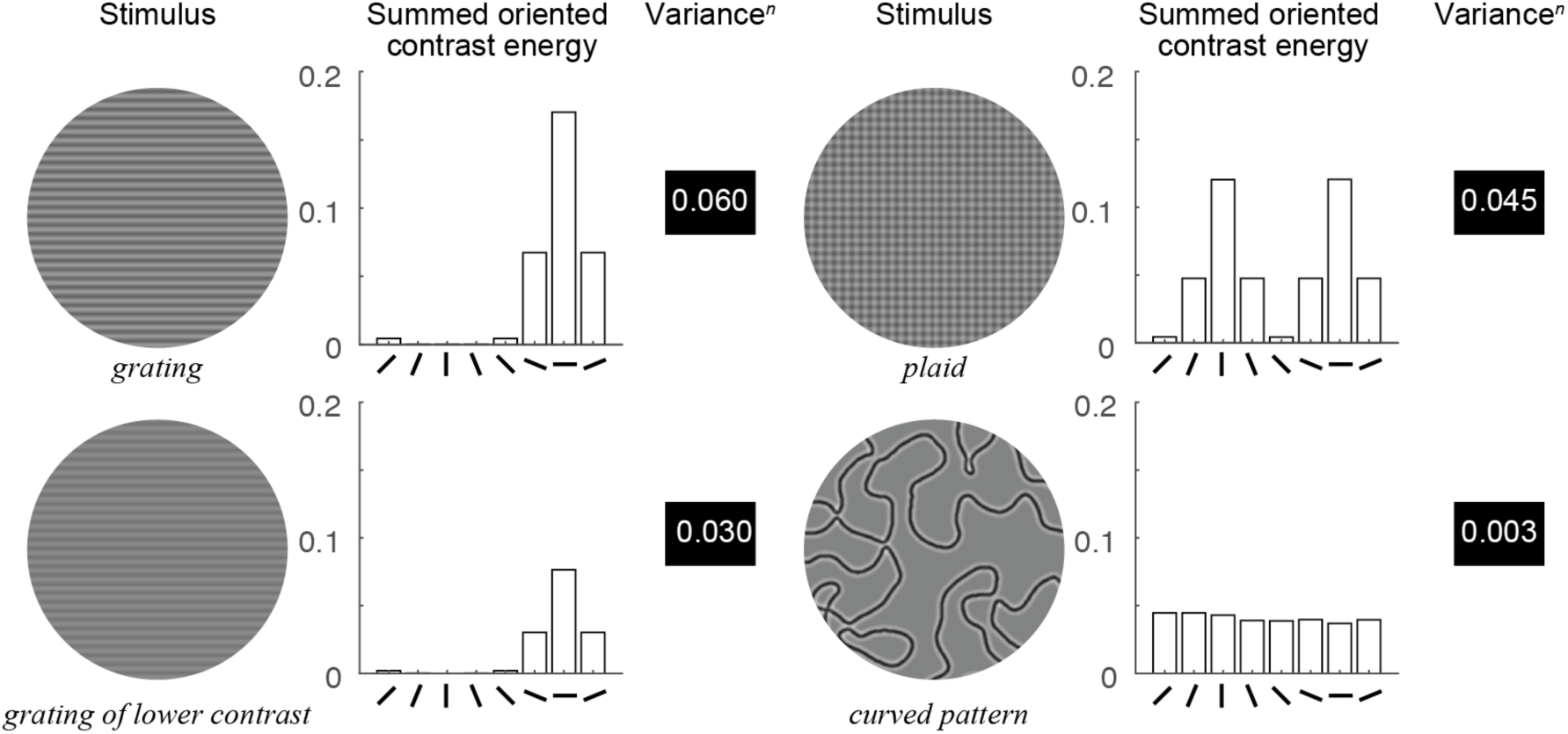
Behavior of the OV model. For each example stimulus, the contrast energy within each of 8 orientation bands is summed across the image (bar plots), which simulates the response for a large receptive field spanning the whole image. The variance*n* (with *n* = 0.5) across these 8 values, monotonically related to the output of the model, is displayed next to the bar plots. This value increases with stimulus contrast (upper left versus lower left) and increases with sparsity of orientations: the high-contrast grating with few orientations present (upper left) has a higher output than the plaid with several orientations present (upper right), which in turn has a higher output than the curved pattern with many orientations (lower right).

### Gamma responses are well predicted by a model that is sensitive to variation in orientation content

We tested how well the OV model explains narrowband gamma responses across the 86 images. For all electrodes, gamma responses across the 86 stimuli are largest for the high contrast grating stimuli of different orientations (Figure 7B, stimuli 39-46 and Supplemental Figures S8 and S9). Gamma is much lower for gratings of reduced contrast (47-50) and plaid patterns (51-55), and is especially weak for circular patterns (stimuli 55-58), with many orientations.

**Figure 7.**
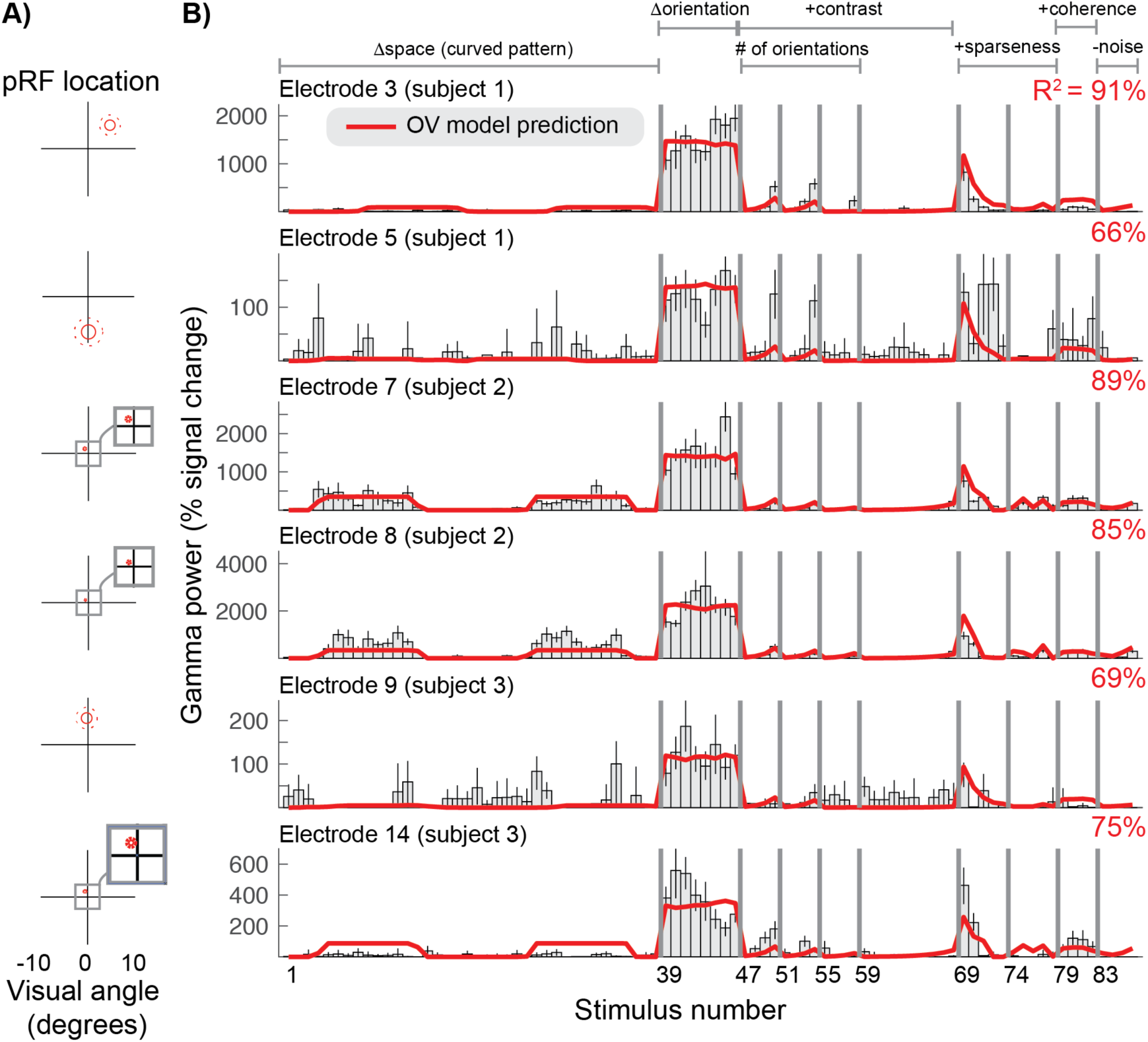
Orientation Variance (OV) model predicts selectivity of gamma responses. **A)** The population receptive field for each electrode was defined by a Gaussian, indicated by the 1-and 2-sd contours (solid and dotted red lines). **B)** The gamma power in percent signal change for six of the 15 electrodes (rows) for all 86 stimuli. Error bars display the 68% range for the bootstrapped responses. The cross-validated predictions of the OV model and overall variance explained (R2) are shown in red.

The OV model has five parameters: the location and size of the pRF (*x, y* and *s*), a gain factor (*g*) and an exponent (*n*). The pRF center and size were derived from separate data, as they could not be robustly obtained by fitting a model to the gamma responses across the 86 images. This is because the stimuli that varied systematically in spatial position (images 1 to 38) induced very little gamma response. We fixed the *x* and *y* position of the population receptive field based on the SOC fits to the broadband data (Figure 4). The sigma parameter was derived from the center parameters (*x, y*) based on an assumed linear relationship between pRF size and eccentricity as reported in (Kay et al., 2013a) (Figure 7A).

Therefore, the only free parameters in fitting the OV model to the gamma responses were the exponent (*n*) and the gain (*g*). We evaluated a range of values for *n* ({.1 .2 .3 .4 .5 .6 .7 .8 .9 1}) and directly fit the gain, and used a leave-one-out cross-validation scheme in order to obtain unbiased estimates of model accuracy. Across electrodes an exponent of *n* = 0.5 predicted most variance in the left-out data, and results with this exponent are reported throughout this paper. Overall, the OV model accounted for the pattern of gamma responses well, with an average of 80% cross-validated variance explained across electrodes.

### Grating-like features in the pRF strongly drive gamma oscillations

The OV model is sensitive to the distribution of contrast energy across orientations within the pRF, but not to image structure remote from the pRF. Because different electrodes have different pRFs, the output of the model can differ between electrodes in response to the same image, and between exemplars of images taken from the same stimulus class (e.g. natural images) for the same electrode.

To illustrate the importance of taking into account the specific pRF location associated with an electrode, we consider several examples. For a stimulus whose orientation changes over space (Figure 8A, left), a large pRF is likely to sample a wide range of orientations. As a result, the spatially summed outputs of different orientation bands are similar, and the variation across these outputs is low. For this reason, for an electrode with a large pRF, the predicted response to the slowly curving pattern (stimulus 10) is smaller than the predicted response to a grating (stimulus 50) (Figure 8B left). This prediction is borne out by the data: gamma is much smaller for the curved patterns compared to the gratings for this electrode (Figure 8B, left panel). In contrast, a very small pRF, such as in foveal areas of visual cortex (electrode 8), is likely to be exposed to a single dominant orientation, even when the full image contains many orientations. As a result, for a small pRF, the OV prediction is similar for the curved patterns and a grating stimulus (Figure 8B, right panel). This example illustrates that it is critical to consider the precise receptive field location of an electrode when investigating gamma oscillations.

**Figure 8.**
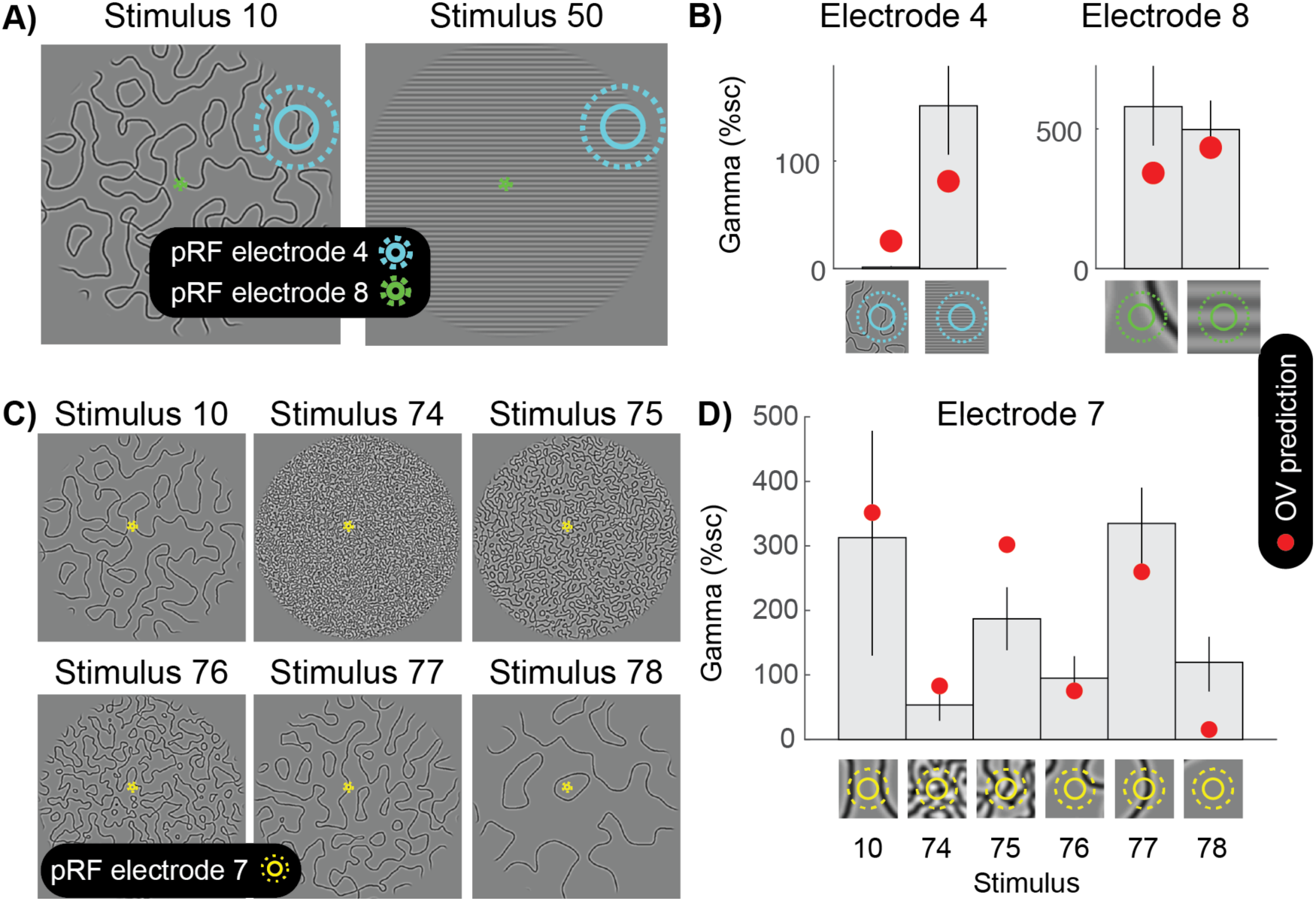
Grating-like features in the pRF strongly drive gamma oscillations. **A)** The population receptive fields (pRF) of two electrodes are overlaid on two different curved patterns (stimulus 10 and 50). Electrode 4 has a large pRF, while electrode 8 has a very small pRF. **B)** The left panel shows that for electrode 4, the OV model predicts a small response for the curved lines and a large gamma response for the grating pattern (red dots). As predicted, there is a small gamma response for the curved lines and a large response for the grating stimulus. Blue circles on the bottom zoom into the stimulus in the pRF. The right panel shows that for electrode 8, the OV model predicts a similar response for the image with curved lines and the image with a grating. As predicted, there is a large gamma response for the curved lines and a large response for the grating stimulus. Green circles on the bottom zoom into the stimulus in the pRF and this shows that from the curved lines, only a relatively straight line falls in the pRF. **C)** Six different images with curves differing in sparseness (stimulus 10, 74, 75, 75, 76, 77 and 78). The population receptive field (pRF) of electrode 7 is overlaid with one and two standard deviations (solid yellow and dotted yellow). **D)** The OV model predicts the largest response when a grating-like feature hits the pRF (red dots). As predicted, the largest gamma responses are observed when the pRF contains grating-like features. Yellow circles on the bottom zoom into the pRF content of each of the stimuli. Error bars display the 68% confidence interval (across bootstraps), and the close up of image in the pRF show the outline of the pRF at 1 and 2 standard deviations (straight and dashed).

The importance of precise pRF locations can also be appreciated by considering responses of a foveal electrode to a variety of curved patterns (Figure 8C-D). When the stimulus is relatively sparse, the small pRF may be exposed to a single dominant orientation (stimuli 10 and 77), resulting in large predicted responses, or no contrast at all (stimuli 76 and 78), resulting in little response. When the stimulus is very dense (stimulus 74), the pRF is likely to be exposed to many orientations, resulting in relatively weak responses. This general pattern of responses is observed in the data, demonstrating that it is critical to take into account the specific pRF location for a visual electrode in order to understand the nature of gamma oscillations.

## Discussion

### Stimulus selectivity of gamma oscillations in visual cortex and their relevance for neural function

The most salient observation is that the narrowband gamma responses were much sparser than the broadband responses. The gamma responses were large for only a few stimuli tested (high contrast gratings), and relatively small or even within the experimental noise for most other stimuli (e.g., noise patterns and textures with a lot of curvature). The sparseness in the narrowband gamma responses was not due to a lack of image contrast or measurement sensitivity. Many of the stimuli that elicited little measured narrowband gamma responses nonetheless elicited large broadband responses. In fact, large broadband responses were observed for most stimuli tested, including all stimuli with high contrast within the electrode’s receptive field (pRF). The much greater sparseness in the narrowband gamma compared to broadband responses was a highly consistent finding, observed in every electrode studied (n=15, coming from 3 participants and spanning V1-V3).

The sparseness in the narrowband gamma responses is striking not just in comparison to the broadband responses reported here, but also in comparison to fMRI responses to the same stimuli measured previously (Kay et al., 2013a). Like broadband, the fMRI response was large for all stimuli with high contrast in the pRF and tended to be larger for curved patterns than for gratings. The similarity in stimulus selectivity between fMRI and broadband is consistent with a number of other studies spanning multiple cortical areas and stimulus manipulations, including motor cortex and finger movements (Hermes et al., 2012), auditory cortex and natural movies (Mukamel et al., 2005), ventral temporal cortex and category selectivity (Jacques et al., 2016), occipital cortex and spatial summation (Winawer et al., 2013), and occipital cortex and pattern selectivity (Hermes et al., 2017b).

The stimulus selectivity of narrowband gamma raises questions about the potential functions of this signal. Gamma oscillations have been hypothesized to play a wide range of cognitive and neural functions, including perceptual binding of visual features (Eckhorn et al., 1988; Gray et al., 1989), prioritizing communication of certain visual information over other information (Fries, 2005), and visual awareness (Engel and Singer, 2001). It is not clear why such functions would be specifically restricted to oriented gratings compared to curved patterns or stimuli with multiple orientations. At high contrast, all of these stimuli are easily visible and all of them elicit robust signals in visual cortex as measured with fMRI and broadband ECoG. Moreover, these stimuli would be expected to elicit strong multiunit responses, given typical models of action potentials in neuronal populations in primary visual cortex (e.g., Rust et al, 2005; Carandini et al, 2005). Finally, it is reasonable to suppose that all of the stimuli require transmission to downstream visual areas in support of recognition and behavior. Narrowband gamma oscillations appear to be a conspicuous outlier among these neuronal signals. The large attenuation in this signal for a variety of simple, high-contrast stimuli suggests that gamma oscillations are unlikely to be a general mechanism for binding, awareness, or information transfer to higher visual areas, and suggests the importance of testing cognitive theories on a variety of different stimulus types. Such theories may hold for some classes of stimuli but not others.

### The orientation-variance (OV) model of gamma responses

Because of the similarity in stimulus selectivity between broadband and fMRI, the same model (SOC) was appropriate to explain both measurements (Kay et al., 2013a). The strikingly different stimulus selectivity of the narrowband gamma responses motivated us to develop a novel image-computable model, the Orientation-Variance (OV) model. The OV model is sensitive to the variation across the spatially pooled outputs of the orientation channels in the population receptive field. The two models share a common first stage, in which contrast energy is computed. As a result, both models predict that response amplitudes should increase with stimulus contrast, in agreement with the observation that fMRI, broadband, and narrowband ECoG responses all increase with contrast. The subsequent calculations of the two models differ from each other, particularly in the order of operations. The SOC model first sums across orientation at each spatial location and then computes variance across space. The OV model does the opposite, first summing across space (within each orientation band) and then computing variance across orientations. The difference in the order of operations has a large effect on the image properties that are emphasized by the model. The SOC model penalizes images (lowers the response) when contrast is high everywhere, akin to surround suppression. Because the OV model first sums across space, responses grow with stimulus size, as is observed for gamma oscillations but not spiking (Gieselmann and Thiele, 2008; Jia et al., 2013; Ray and Maunsell, 2011; Self et al., 2016) or BOLD (Press et al., 2001; Zenger-Landolt and Heeger, 2003; Zuiderbaan et al., 2012).

For both models, the computations are made within the population receptive field. Therefore, the question of whether gamma is induced by an image, like the question of whether any neural signal is induced by an image, is only sensible if we take into account the *specific receptive field location* of the neurons or neuronal population under consideration, as shown in Figure 8.

The distinction between the broadband and gamma responses, and the need for different models, was emphasized by choices we made in selecting stimuli and in analysis of the data. For example, had we only used gratings of varying contrast, the gamma, broadband, and BOLD signals would have been found to collectively rise together (though perhaps differing in the precise shape of the contrast response functions (Henrie and Shapley, 2005; Lima et al., 2014)). Similarly, when using only grating stimuli which elicit large gamma responses, e.g (Scheeringa et al., 2011), trial-to-trial variance in responses may reflect global factors such as attention or arousal; these global factors might modulate multiple signals, such as gamma power and BOLD, thereby causing the two signals to be correlated within the experimental paradigm. By systematically exploring variation in stimulus properties such as the number of component orientations (gratings vs plaids vs circular patterns), we were able to reveal opposing effects on broadband and gamma responses, similar to the effects of manipulating stimulus size, which has opposite effects on the level of broadband and gamma responses (Ray and Maunsell, 2011). These different response patterns show the importance of testing a wide range of stimuli. In addition, the method of separating the ECoG measurement into two components was also important. Had we simply characterized the gamma response as the band-limited power increase over baseline, the gamma responses would have appeared less sparse. For example, the noise patterns in Figure 1 cause power increases in the gamma band (∼30-80 Hz) for two of the three electrodes. However, because the power increase spans higher frequencies as well and contains no clear peak, the response is better characterized as a broadband response, rather than as a narrowband gamma oscillation (Lopes da Silva, 2013). This is reflected in how we compute the two signals. We do not assume a direct relationship to temporal frequency bands since the two signals can overlap in their spectra; rather, we take a model-based approach in which we separate out a peaked response (oscillatory) from one that is non-peaked (broadband). This approach is motivated by the fact that the two responses can be modulated independently (e.g., the peaked response decreases with the number of orientations, whereas the broadband response increases). Additionally, independent modulation of a broadband component spanning frequencies from 1 to 200 Hz has been previously demonstrated by principal components analysis of spectral power (Miller et al., 2009a).

To better understand gamma oscillations in the context of natural vision, we computed the outputs of the OV and SOC models to a large collection of natural images (Olmos and Kingdom, 2004). These computations reveal a few interesting patterns. First, the outputs of the two models show some degree of positive correlation, consistent with the fact that both model outputs increase with stimulus contrast within the pRF (Figure 9). Second, the responses to gratings are distinct from the responses to natural images, especially for models with larger pRFs (upper panel). This is because the OV output to gratings is unusually high compared to natural images, whereas the SOC output is within or close to the range of outputs to natural images. These patterns are generally consistent across models fit to the 15 electrodes we tested (Supplementary Figure S10). High-contrast gratings are often used in studies of gamma oscillations, e.g. (Eckhorn et al., 1988; Hoogenboom et al., 2006; Rohenkohl et al., 2018); using these stimuli as a benchmark, we find that the OV outputs for natural images are, in general, strikingly low. However, natural images occasionally have oriented high-contrast energy within the pRF and these give rise to large OV output (Figure 9). Hence, the image properties to which the OV model is sensitive exist in natural images, but in far lower quantities than they do for oriented gratings (and other stimuli such as bars (Gray and Singer, 1989; Gray et al., 1989) and circular gratings (Hoogenboom et al., 2006) that are locally similar to oriented gratings). Note that the range of the OV outputs for these natural images (on the order of ∼100%) is comparable to the gamma band responses measured in monkey ECoG to gray-scale natural images reported by Brunet et al (Brunet et al., 2015). To the degree that the OV model accurately predicts narrowband gamma responses and the SOC model accurately predicts broadband responses, these results indicate that for most natural gray-scale images (with occasional exceptions), relatively low levels of gamma oscillations are expected.

**Figure 9.**
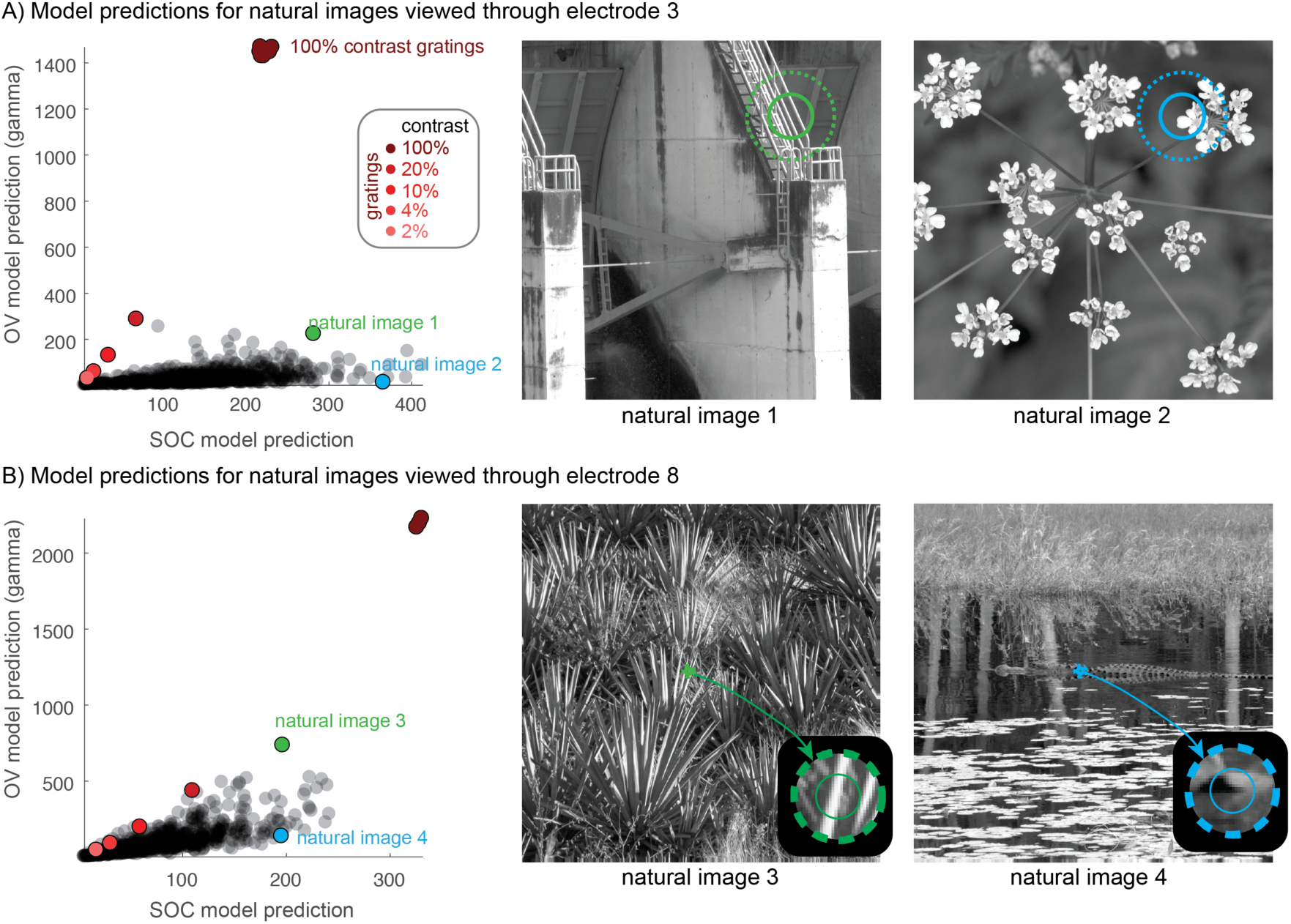
OV and SOC model predictions for images of natural scenes. **A)** The OV and SOC outputs are plotted for a set of gray-scale photographs of scenes, with model parameters from electrode 3. The units are in percent signal change, as in Figs 4 and 7. Each gray dot is the output of the two models for one image. The red dots are the model outputs for grating stimuli of varying contrast. The cluster of red dots at 100% contrast displays high-contrast gratings of different orientations (stimuli 39-46). The green and blue dots correspond to two images with large OV and SOC outputs, respectively. The right panels show these two images with the electrode pRF location superimposed (1 and 2 SDs). Natural image 1, with a high OV output, has image features in the pRF that look like a grating. The OV output to images of natural scenes are much lower than the responses to high contrast gratings. **B)** Same as panel A, but for electrode 8, including a zoom into the pRF location.

Finally, we note that the absolute scale of the responses differs substantially between the models, with a larger maximal response for the OV model, especially for gratings (for example, up to 2000% signal increase over baseline compared to 300% for the SOC model; Figure 9 bottom). This difference in amplitude exists not only in the model outputs, but also in the data (e.g., Figures 2, 3, 4 and 7). Why are gamma responses so much larger than broadband responses? Broadband responses are thought to arise from asynchronous neural activity (Miller et al., 2009b), which results in substantial cancellation in the pooled field potential (Butler et al., 2017; Hermes et al., 2017b; Krusienski et al., 2011; Winawer et al., 2013). Narrowband gamma responses, in contrast, are thought to reflect synchronous activity across a neuronal population. The synchronous response, even if it comes from a much smaller neuronal population, can result in a much larger macroscopic field potential (Butler et al., 2017; Hermes et al., 2017b; Winawer et al., 2013). This likely explains why gamma oscillations, even when the percentage signal change is quite large, can show little correlation with the BOLD signal (Hermes et al., 2017b) or multiunit activity (Jia et al., 2013; Ray and Maunsell, 2011). Hence the large size of the field potential generated by gamma oscillations does not imply a high level of neuronal population activity in terms of energy demand or spike rates. In contrast, substantial increases in broadband power do generally correlate with both high energy consumption and high firing rates (Manning et al., 2009; Ojemann et al., 2013; Ray and Maunsell, 2011) (though for an exception see (Leszczynski et al., 2019)).

For this study, we did not model or explore the effect of color on gamma oscillations, a feature which has recently been shown to strongly modulate gamma (Peter et al.; Shirhatti and Ray, 2018). Future work will be required to link our models of spatial pattern with models of chromatic sensitivity.

### Gamma oscillations and gain control

Why do some specific stimuli elicit gamma responses? Both in this report and elsewhere (Gieselmann and Thiele, 2008; Henrie and Shapley, 2005; Jia et al., 2013; Ray and Maunsell, 2011), large-amplitude gamma oscillations are found for stimuli that are high in contrast, spatially extended, and with few orientations. These three image properties—contrast, spatial extent, and limited orientations—all produce larger outputs in the OV model, and interestingly, are all associated with gain control or suppression in neuronal circuits.

First, stimulus contrast has been linked to inhibition in divisive normalization models of primary visual cortex (Heeger, 1992). According to this model, gain control increases with local stimulus contrast, possibly via shunting inhibition (an increase in membrane conductance) or a reduction in recurrent amplification (Sato et al., 2016). Although neuronal responses such as spike rates tend to increase with contrast, the rate of increase is slower at higher contrast (Albrecht and Hamilton, 1982), consistent with mechanisms of increasing gain control at higher contrast (Albrecht and Geisler, 1991; Heeger, 1992). Second, stimulus extent is linked to suppression in that larger stimuli stimulate the inhibitory surrounds of neuronal receptive fields, thereby reducing the neuronal response (Allman et al., 1985). Third, for a large stimulus, surround suppression is more effective when an annulus and a central stimulus match in orientation (Cavanaugh et al., 2002; DeAngelis et al., 1994; Knierim and van Essen, 1992).

In summary, each of these three stimulus properties (high contrast, large spatial extent, limited orientations) is associated with more gain control or suppression as well as a larger OV output, consistent with the proposal that gamma oscillations are a biomarker of gain control or normalization (Ray et al., 2013). In addition to the patterns observed here, this proposal has been supported by recordings in macaque MT: when a null-motion stimulus is added to a preferred motion stimulus, spike rates decrease, indicating an inhibitory effect of the null motion stimulus, and gamma oscillations increase in amplitude (Ray et al., 2013). This again shows that a stimulus configuration that increases inhibition also increases the amplitude of gamma oscillations. We note that the link between gamma oscillations and gain control does not necessarily indicate what causal role, if any, the oscillations have in neural processing. At a minimum, the oscillations may serve as a biomarker of gain control circuits, useful to the experimenter but not necessarily to the organism producing them. Whether or not the oscillations are critical for implementing gain control requires further study.

One exception to the link between inhibition and gamma power in this study is the effect of the number of component orientations: increasing the number of superimposed gratings decreases the OV output and the power of narrowband gamma (Figure 3), yet causes an increase in cross-orientation suppression in visual cortex (Bonds, 1989; Concetta Morrone, M. et al., 1982). This breaks the pattern by which stimulus properties that cause more inhibition also cause larger-amplitude gamma oscillations. Cross-orientation suppression and surround suppression differ in several ways, including in their temporal properties, with surround suppression slightly delayed (Smith et al., 2006). This supports the possible interpretation that cross-orientation suppression is inherited from earlier processing in a feedforward manner, whereas surround suppression depends on intra-cortical connections (either within a cortical area and/or via feedback). A large increase in gamma oscillations may therefore reflect locally implemented inhibition, as in surround suppression, but not inherited suppression, as in cross-orientation suppression. More generally, the fact that these two types of suppression likely have different underlying mechanisms, and different effects on gamma oscillations, highlights the importance of considering the circuit-level implementation of computations such as suppression and gain control.

At a more abstract level, the same image properties that are associated with gain control can also be described as signatures of image redundancy (or predictability). Since gamma oscillations tend to increase in the presence of gain control, they can also be described as increasing in the presence of image redundancy or predictability (Vinck and Bosman, 2016). In fact, early descriptions of center-surround visual receptive fields proposed that the surround suppression was part of a coding strategy to compress signals elicited by natural images which contain a lot of redundancy in the form of spatial correlations (Barlow, 1961).

### Gamma oscillations and neuronal circuits

While there is growing evidence that gamma oscillations increase in the presence of gain control or inhibition, the circuitry underlying these inhibitory mechanisms are not firmly established. For example, contrast gain control modeled as divisive normalization might be implemented biologically as shunting inhibition (synaptic inhibition which changes the neuronal membrane conductance) (Carandini and Heeger, 1994; Carandini et al., 1997). In this implementation, the notion of inhibition in the model (signal reduction by division) is literally an increase in inhibitory neural signals. Alternatively, normalization could be implemented by a circuit that reduces excitation, which in turn also *reduces* inhibition rather than increases it, as is the case for inhibition-stabilized networks (ISNs) (Ozeki et al., 2009; Tsodyks et al., 1997). In ISNs, the un-normalized state (e.g., low contrast, no suppressive surround) has a high level of recurrent excitation and inhibition, with the inhibition serving to stabilize the network. Stimulus manipulations that result in an increase in gain control, such as the addition of a surrounding stimulus or an increase in contrast, paradoxically result in a withdrawal of inhibition. This, in turn, destabilizes the network, allowing the activity to either die off or explode. Gamma oscillations may be more likely to arise (or to increase in amplitude) in this destabilized state. In the extreme, large, high-contrast oriented gratings can even trigger seizures in patients with photosensitive epilepsy or cause discomfort in healthy subjects (Harding et al., 2005; Hermes et al., 2017a; Wilkins et al., 1984).

A large number of studies have tried to explain the circuit mechanisms that underlie gamma oscillations, including via explicit computational models (e.g., Ainsworth et al., 2012; Buzsáki and Wang, 2012; Womelsdorf et al., 2014). Some of these models predict modulations in the amplitude of gamma oscillations as a function of labeled stimulus properties, such as size or contrast (Jia et al., 2013). Yet to our knowledge, none of these models operates on visual inputs in the sense of arbitrary pixel values, as does our image-computable OV model. Our model, however, does not describe the neuronal circuitry that produces the oscillations. A more complete understanding of this intensely studied neural signal will likely require a unified account that both generalizes to arbitrary images and also specifies the circuitry that underlies oscillations.

## Conclusions

Gamma oscillations in human visual cortex are elicited by distinct types of visual inputs compared to fMRI BOLD and ECoG broadband responses. We developed an image-computable ‘orientation-variance’ model, which accounts for the amplitude of gamma oscillations across many stimuli. In this model, gamma oscillations are driven by increases in contrast and by variance across orientation channels in the population receptive field. These findings suggest that gamma oscillations reflect circumstances in which neural circuits exhibit strong normalization or gain control.

## Methods

### Ethics statement and subjects

ECoG data were recorded in three subjects (mean age 28, 2 women) who had electrodes implanted for the clinical purpose of epilepsy monitoring. Subjects gave informed consent and the study was approved by the Stanford University IRB and the ethics committee at the University Medical Center Utrecht in accordance with the 2013 provisions of the Declaration of Helsinki.

### Stimuli and task

Static visual images were viewed from a distance of ∼50 cm and spanned approximately 20 degrees of visual angle. Images were presented for 500 ms, followed by a grey screen for 500 ms. There were 86 different gray scale images that contain spatial frequencies of 3 cycles per degree. Images included curved lines with varying apertures spanning parts of the visual field, full screen gratings varying in orientation, plaids, circular patterns with 16 orientations and curved lines varying in contrast, gratings and curved lines varying in sparseness, gratings varying in coherence and curved lines with different levels of noise. These stimuli could roughly be grouped into the categories of SPACE (1 to 38), ORIENTATION (39 to 46), CONTRAST (47 to 68), SPARSITY (69 to 78), and COHERENCE (79 to 86) All images are shown in the Supplemental Materials Figures S1-S5. Images were created in similar manner as in (Kay et al., 2013a), with only images 79-86 being a new category. Each image was repeated several times (subject 1: 15 times, subject 2: 9 times, subject 3: 12 times).

### ECoG procedure

ECoG electrodes were placed on the left hemisphere in subject 1 and on the right hemisphere in subjects 2 and 3. ECoG data were recorded at 1528 Hz through a 128-channel Tucker Davis Technologies recording system (http://www.tdt.com) (subjects 1 and 2) and at 2048 Hz through a 128-channel Micromed recording system (subject 3). To localize electrodes, a computed tomography (CT) scan was acquired after electrode implantation and co-registered with a preoperative structural MRI scan. Electrodes were localized from the CT scan and co-registered to the MRI, and positions were corrected for the post-implantation brain shift (Hermes et al., 2010). Electrodes that showed large artifacts or showed epileptic activity, as determined by the patient’s neurologist were excluded, resulting in 116/107/54 electrodes with a clean signal. Offline, data were re-referenced to the common average, low pass filtered and the 1528 Hz data were resampled at 1000 Hz for computational purposes using the Matlab resample function. Line noise was removed at 60, 120 and 180 Hz (Stanford) using a third order Butterworth filter, data from UMC Utrecht did not contain much line noise and were not filtered. In further analyses, we only included electrodes that were located on visual areas V1, V2 or V3 with population receptive fields that were within the stimulus (∼10 degrees from the fovea).

### ECoG analyses

#### Spectral analysis

We calculated power spectra and separated ECoG responses into broadband and narrowband gamma spectral power increases in similar manner as before (Hermes et al., 2015). For each stimulus condition, the average power spectral density was calculated every 1 Hz by Welch’s method (Welch, 1967) with a 200 ms window (0 – 500 ms after stimulus onset, 100 ms overlap). A Hann window was used to attenuate edge effects. ECoG data are known to obey a power law and to capture broadband and narrowband gamma increases separately the following function (*F*) was fitted to the average log spectrum from 30 to 200 Hz (leaving out 60 Hz line noise and harmonics) from each stimulus condition:

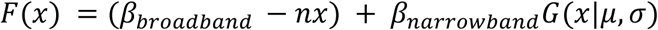

In which,

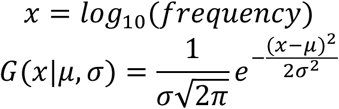

with 0.03 < σ < 0.08 and 30*H*_*Z*_ < 10^*μ*^ < 80*H*_*Z*_.

The slope of the log-log spectral power function (*n*) was fixed for each electrode by fitting it based on the average power spectrum of the baseline. Confidence intervals were estimated using a bootstrap procedure: for each stimulus condition C with N_c_ trials, N_c_ trials were drawn randomly with replacement and power spectra were averaged. The function *F* was fit to the average log_10_ power spectrum from these trials and the β parameters were estimated. This was repeated 100 times, resulting in a distribution of broadband and narrowband weights.

The β parameters in function *F(x)* have units of log_10_ power and from this we derived the percent signal change in broadband and gamma power. The percent signal change is defined as:

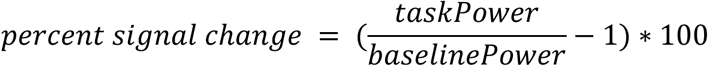

in which,

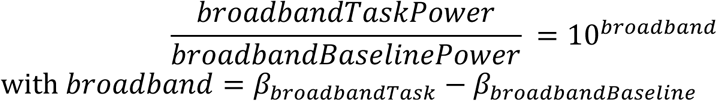

and,

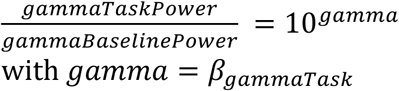

Given the assumption that gamma power is close to zero during the baseline period.

Time frequency analysis was performed around stimulus onset (−500 to 1000 ms) with a multitaper approach (Percival and Walden, 1993) using chronux (http://www.chronux.org/ (Mitra and Bokil, 2008)). A moving window of 200 ms (with overlap of 50 ms) and the use of 5 tapers result in a frequency resolution of 5 Hz, with a spectral smoothing of ±15 Hz. To normalize the responses to baseline, the average spectrum from all inter trial intervals 250–500 ms after stimulus offset was computed and divided from every time bin. The base 10 log was then computed on this normalized power and plotted (Figure 1).

### Model

#### Image preprocessing

Images of 800×800 pixels were downsampled to 240×240 pixels for computational purposes and converted into a contrast image: all pixel values between 0 and 254 were rescaled to a range from zero to one and the background luminance was subtracted (0.5), resulting in all pixel values in a range from −0.5 to 0.5 with the background corresponding to zero. Images were zero padded with 15 pixels on each side to reduce edge effects, resulting in images of size 270×270.

#### Oriented contrast energy (step 1)

After preprocessing, the images were filtered with isotropic Gabor filters with 8 different orientations and 2 quadrature phases covering positions on a grid of 135×135. Since the stimuli were band-pass, filters had one spatial scale with a peak spatial frequency of 3 cycles per degree. The filters were scaled such that the response to a full-contrast optimal grating was 1. After quadrature-phase filtering, the outputs were squared, summed, and square-rooted and the results can be expressed as oriented contrast energy (Figure 5A):

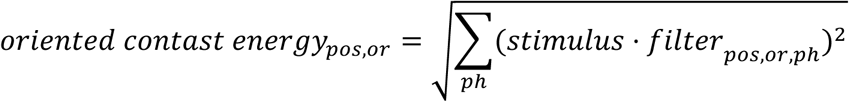

With *pos* the 135×135 positions in the image, *or* the 8 orientations, *ph* the 2 phases of the Gabor filter, and *stimulus* is the preprocessed image and *filter*_*pos, or, ph*_ the oriented Gabor filter with a particular position, orientation and phase.

#### Spatial summation (step 2)

For each orientation, the oriented contrast energy is then summed across space (for each orientation) using isotropic 2D Gaussian weights (Figure 5B):

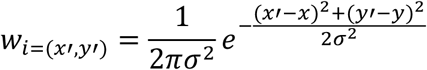

where *w*_*i=(x’,y’)*_ is the weight at position *i* indexed by coordinates *x’* and *y’*; *x* and *y* indicate the center of the Gaussian; and *s* is indicates the standard deviation of the Gaussian. Note that because of the scaling term, the sum of the weights equals one:

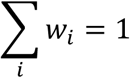

The first two steps result in eight values (one per orientation) for the oriented contrast energy summed within the pRF defined by the Gaussian.

#### Variance (step 3)

A value for the summed oriented contrast energy will be high if an orientation has a high contrast in the area described by the 2D Gaussian and small in case an orientation has low contrast in the area described by the Gaussian. The variance is then calculated across these eight summed oriented contrast energy values, exponentiated with an exponent and multiplied by a gain (Figure 5C):

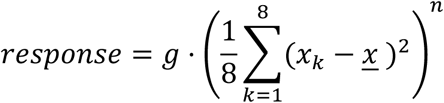

With *x*_*k*_ the summed contrast energy or orientation *k* of which there are a total of 8, *n* an exponent, *g* the gain and <u>*x*</u> the mean summed contrast energy. When one orientation is present and others are not (e.g. [1 0 0 0 0 0 0 0]), the variance will be high. When several orientations are present to different degrees (e.g. [.4 .5 .4 .5 .4 .5 .4 .5]), the variance will be low (Figure 6). The image contrast and the number of orientations drive the response of the OV model, and the model has five parameters: *x, y, σ, g* and *n*.

### Model fitting

The SOC model was fit to the broadband data and the OV model was fit to narrowband gamma data using leave one out cross validation.

#### Fitting broadband power changes with the SOC model

The SOC model was previously developed to explain fMRI signal changes for many of the images that were used here (Kay et al., 2013a). To explain ECoG broadband changes with this model, we used a very similar fitting approach.

We fit the SOC model to ECoG broadband response amplitudes from each electrode. Model fitting was performed using nonlinear optimization (MATLAB Optimization Toolbox) with the objective of minimizing squared error. To guard against local minima, we used a variety of initial seeds for the *c* and *n* parameters. For every combination of *c* and *n*, where *c* is chosen from {.1 .4 .7 .8 .9 .95 1} and *n* is chosen from {.1 .3 .5 .7 .9 1}, we optimized *x, y, σ*, and *g* with *c* and *n* fixed, and then optimized all of these parameters simultaneously. To optimally fit the pRF location, we first seed the pRF in the center of the stimulus and estimated the model parameters from the SPACE stimuli, and then use the estimated *x, y* and *σ* to fit the model again on all stimuli. To get an unbiased estimate of the model accuracy we fit the model using leave one out cross-validation.

#### Fitting narrowband power changes with the new OV model

The OV model has five parameters that need to be estimated: the *x, y*, and *σ* of the Gaussian that define the location and size of the population receptive field, an exponent *n* and a multiplicative gain *g*. The OV model was fit to the ECoG narrowband gamma power changes. The *x* and *y* position of the pRF were derived from the SOC model fit to the broadband data from the same electrode. There is a consistent relationship between the eccentricity of the pRF in V1, V2 and V3 and its size (Kay et al., 2013a), and we used this relationship to calculate the size *σ* of the pRF. The only parameters that are left to be estimated through fitting are the gain *g* and the exponent *n*. We tested an exponent of {.1 .2 .3 .4 .5 .6 .7 .8 .9 1} and derived the gain through a linear regression with least squares between the model output and all data except one stimulus. Model performance was then tested on the left out stimuli (leave one out cross-validation).

#### Model Accuracy

The model performance was evaluated on the data for the left out stimulus (leave one out cross-validation). As a measure for model performance we calculated the coefficient of determination (COD):

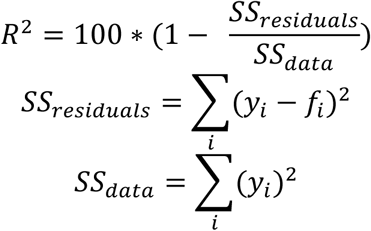

where *y*_*i*_ is the measured response amplitude and *f*_*i*_ is the predicted response amplitude for stimulus *i*. Note that *R*^*2*^ is defined here with respect to zero, rather than with respect to the mean response (similar as in (Kay et al., 2013a)). This metric of prediction accuracy accounts for both accuracy of the mean and the variance across conditions. A model that predicted only the mean (i.e. predicted the same response for each condition) would have, averaged across electrodes, a 24% accuracy for gamma oscillations, much lower than the 80% accuracy of the OV model, and 42% for the broadband response, much lower than the 82% accuracy of the SOC model.

### Natural image simulations

We calculated the predictions of the SOC and OV models for a set of natural images. We used a large collection of 771 natural photographs from the McGill Colour Image Database (Olmos and Kingdom, 2004). These images were converted to grayscale luminance values using supplied calibration information, cropped to square, and downsampled to 240×240 pixels. The images were then further processed and filtered in the exact same way as the stimuli used in the main experiment. Simulated outputs from the SOC and OV models were calculated using the SOC and OV parameters for every electrode. This resulted in 771 simulated SOC and OV responses for each of the 15 electrodes.

## Supporting information

Hermesetal_SupplementalMaterials

## Data availability statement

To foster reproducible research, we make the data and code publicly available via a permanent archive on the Open Science Framework (*url* to be supplied upon acceptance for publication). The data provided conform to the Brain Imaging Data Structure (BIDS) (Gorgolewski et al., 2016; Holdgraf et al.) for ease of use by other researchers.

## Acknowledgements

This work was supported by the Dutch Organization for Scientific Research grant 016.VENI.178.048 to DH and the National Institute of Mental Health grant R01MH111417-01 to JW and NP. The authors thank the Parvizi lab at Stanford, the Stanford Human Intracranial Cognitive Electrophysiology Program (SHICEP), and the Ramsey lab, Cyrille Ferrier and the neurophysiology team at the UMC Utrecht for their help in recording the ECoG data, and we thank David Heeger for helpful discussions and comments on an earlier draft of the manuscript.

