## Supplementary material for "An image-computable model for the stimulus selectivity of gamma oscillations": Hermesetal_SupplementalMaterials

### Supplemental Materials

This document contains the supplemental materials to: *"An image-computable model for the stimulus selectivity of gamma oscillations"*
Dora Hermes, Natalia Petridou, Kendrick Kay, Jonathan Winawer

#### Figure S1

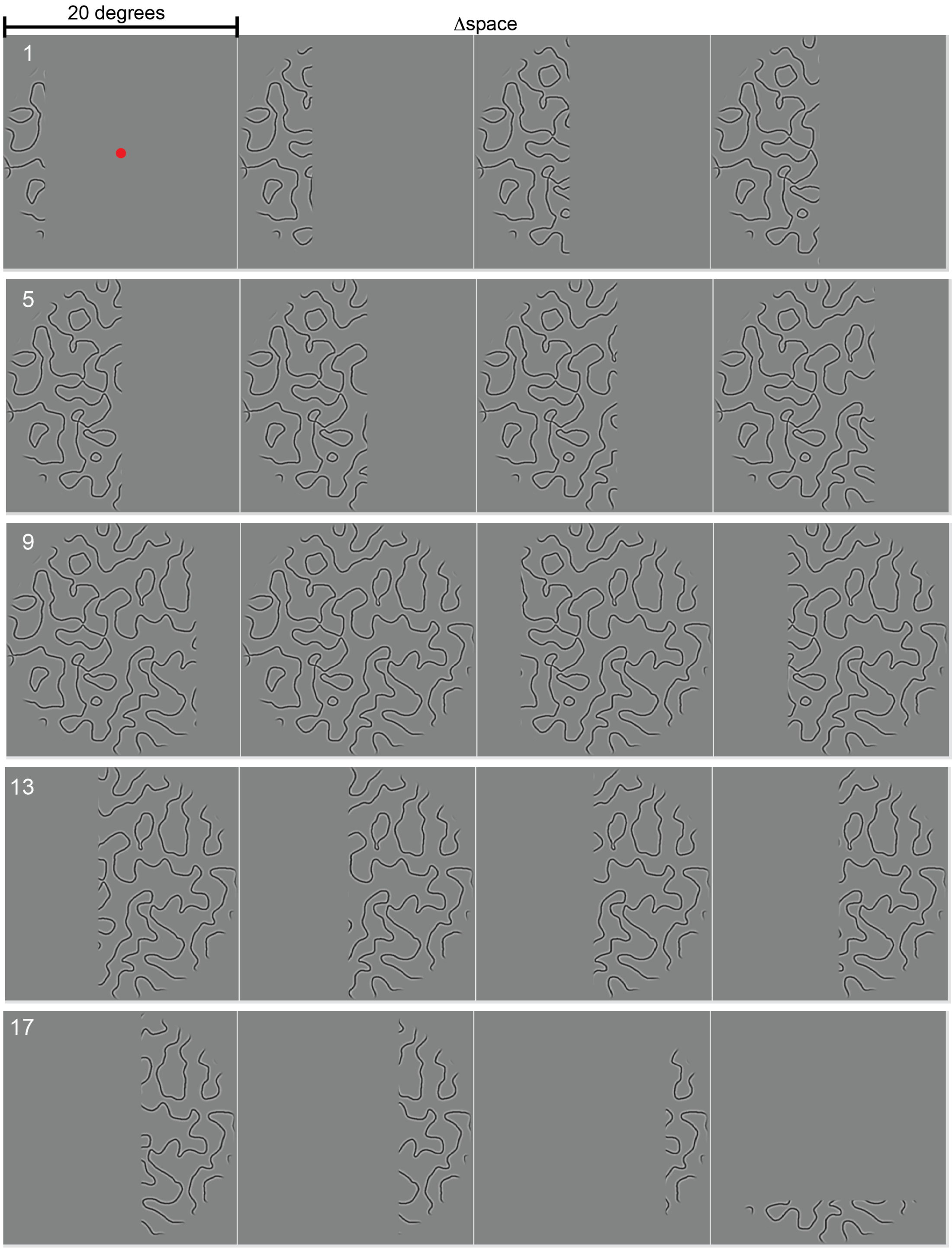

#### Figure S2

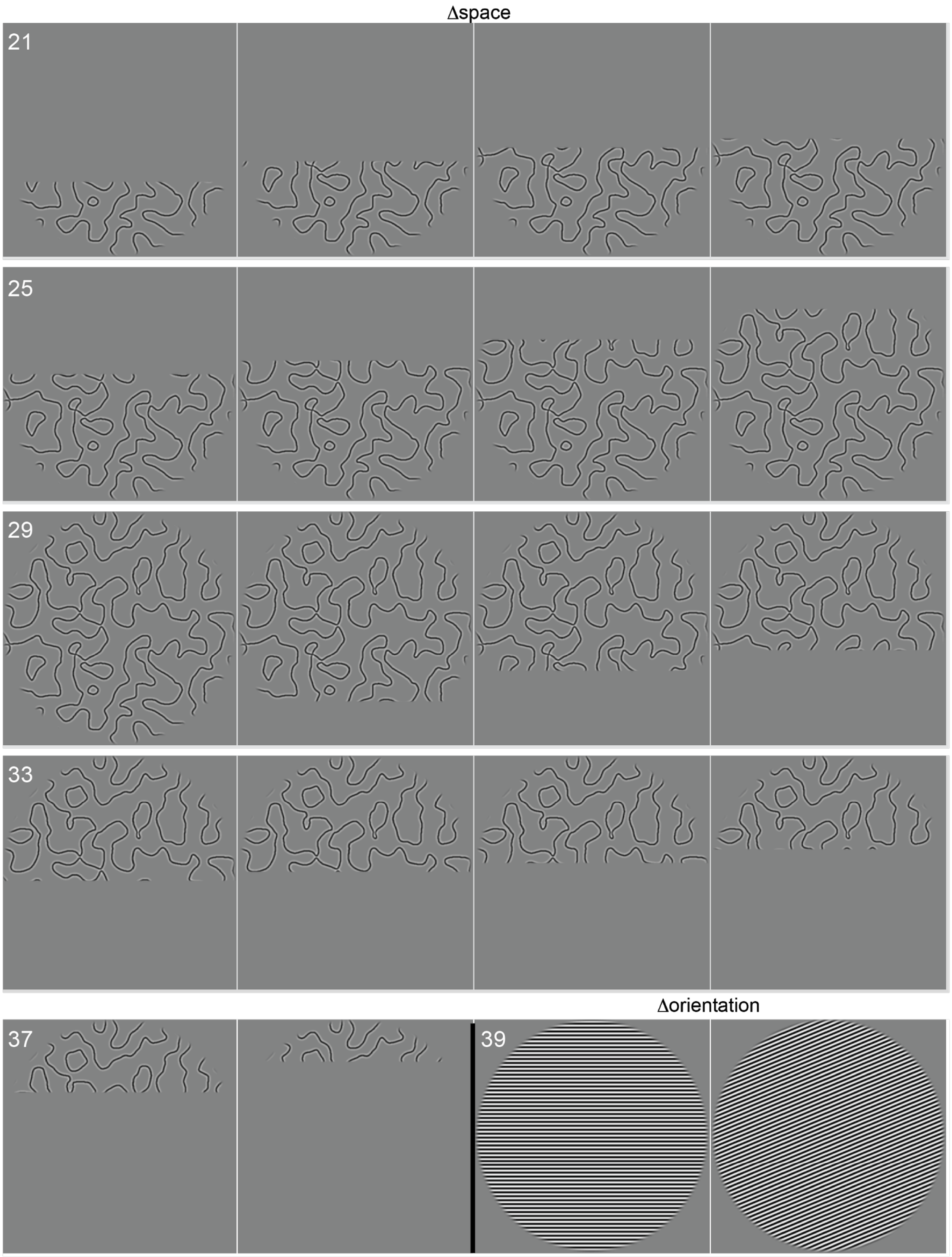

#### Figure S3

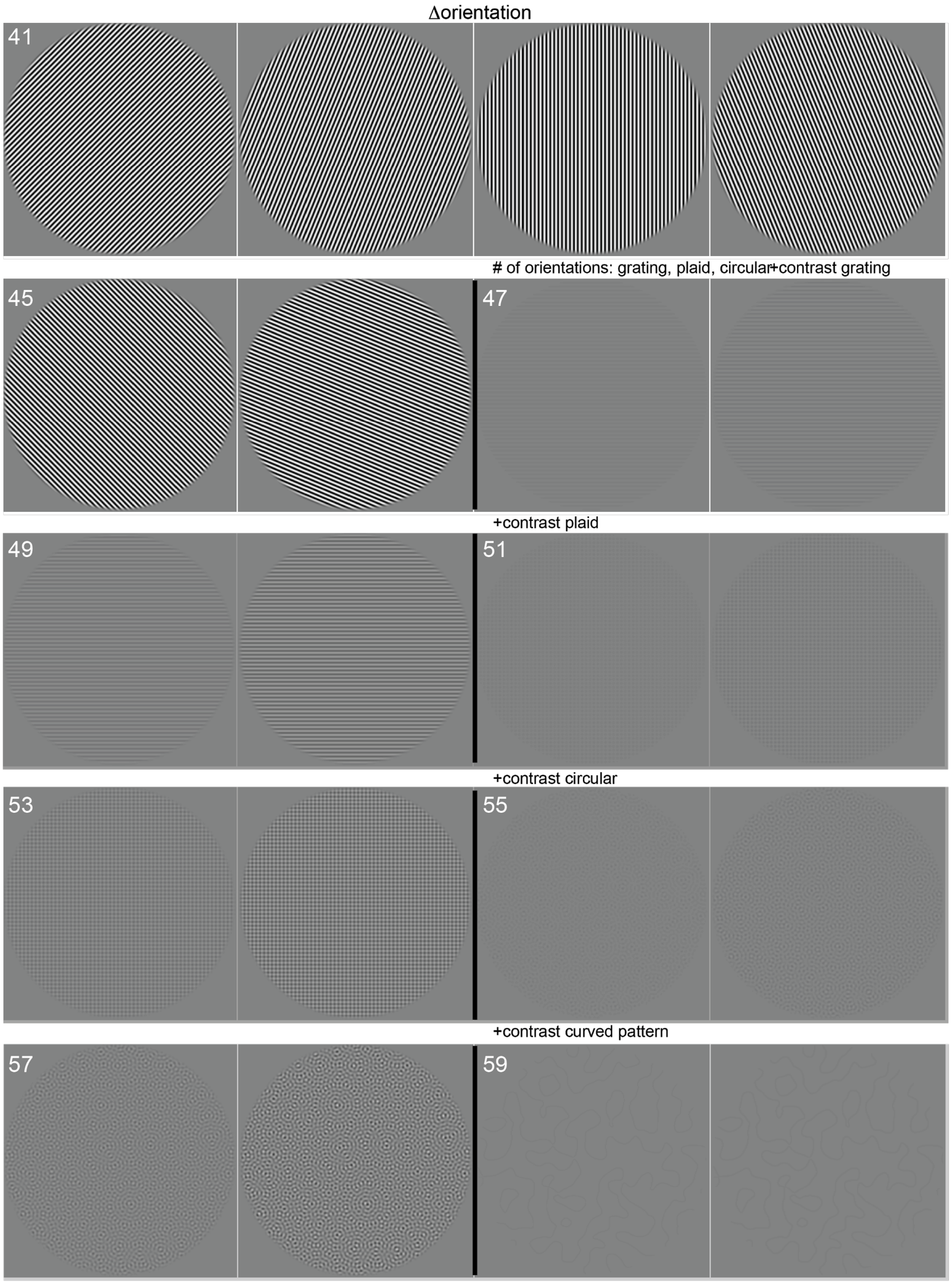

Δorientation
41
### of orientations: grating, plaid, circular+contrast grating
45
47
+contrast plaid
49
51
+contrast circular
53
55
+contrast curved pattern
57
59

#### Figure S4

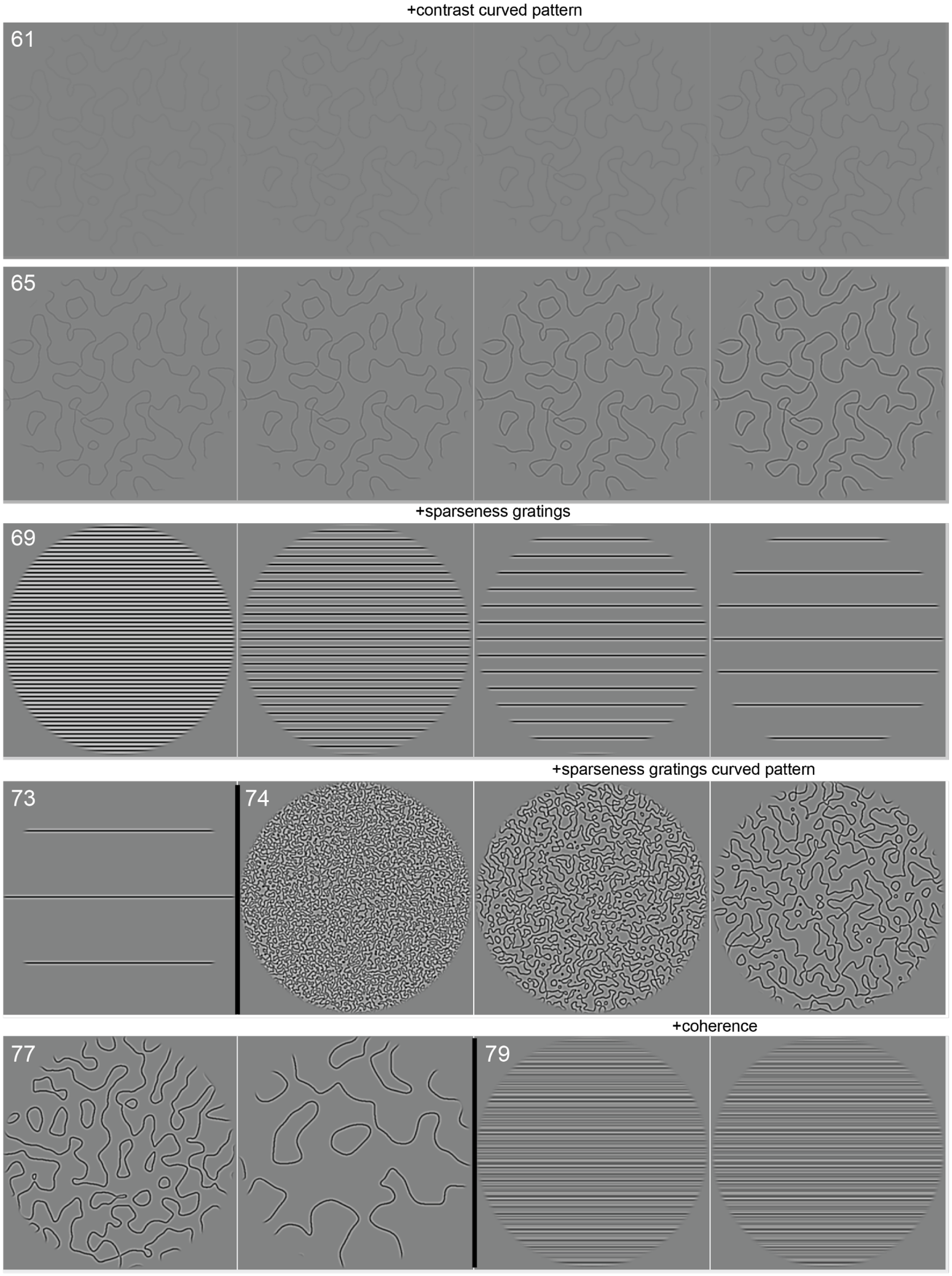
+contrast curved pattern
61
65
+sparseness gratings
69
+sparseness gratings curved pattern
73
74
+coherence
77
79

#### Figure S5

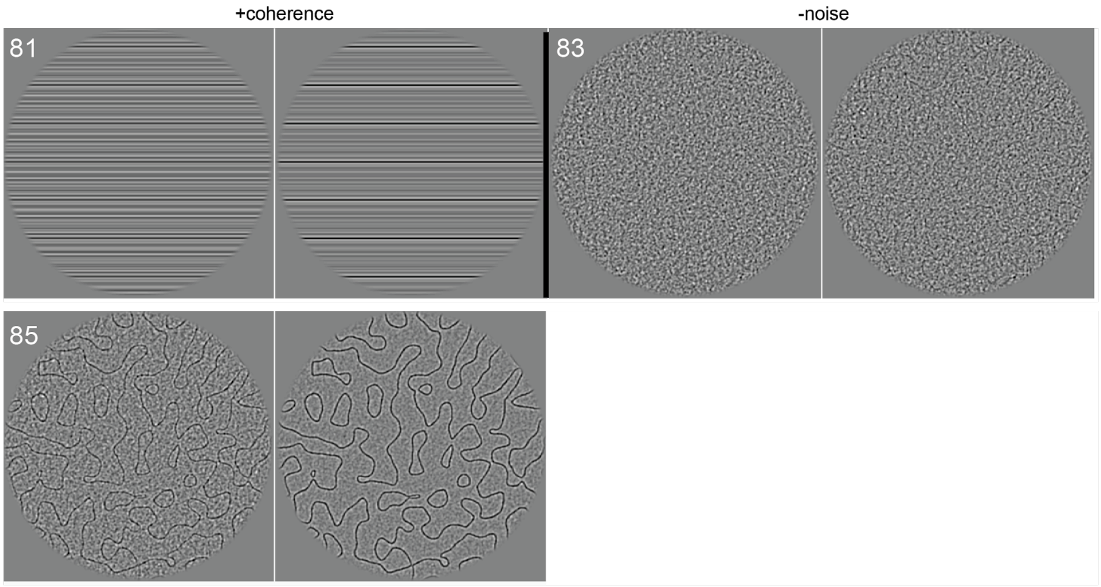


**Figure S1-S5. Stimuli.** These 86 stimuli were shown in random order for 500 ms to the subjects. The top stimulus shows how a fixation dot was presented in the center, and the subjects had to indicate when it changed color. Stimulus 1-38 different in the space that was covered by the contrast pattern. Stimulus 39-46 contained gratings of varying orientation. Stimulus 47-50 contained gratings (1 orientation) of increasing contrast. Stimulus 51-54 contained plaids (2 orientations) of increasing contrast. Stimulus 55-58 contained curved patterns (multiple orientations) of increasing contrast. Stimulus 59-68 contained curved patterns of increasing contrast. Stimulus 69-73 contained gratings of increasing sparseness and stimulus 74-78 contained curved patterns of increasing sparseness. Stimulus 79-82 was a grating increasing in coherence. Stimulus 83-86 contained a curved pattern decreasing in the amount of noise.

#### Figure S6

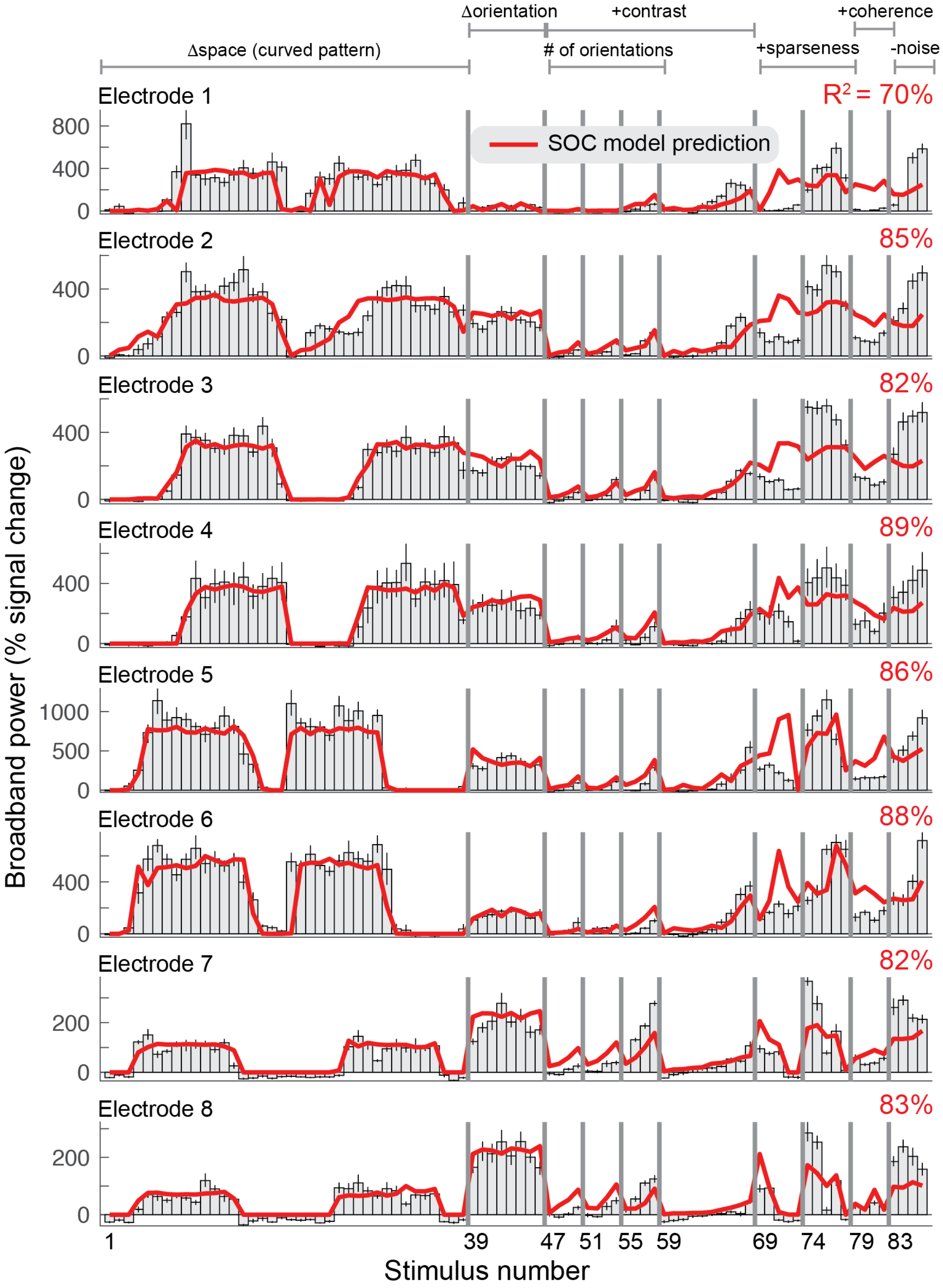

Δorientation
+contrast
+coherence
Δspace (curved pattern)
### of orientations
+sparseness
-noise
Electrode 1
R² = 70%
SOC model prediction
800
400
0
Electrode 2
85%
400
0
Electrode 3
82%
400
0
Electrode 4
89%
400
0
Electrode 5
86%
1000
500
0
Electrode 6
88%
400
0
Electrode 7
82%
200
0
Electrode 8
83%
200
0
Broadband power (% signal change)
1
39
47
51
55
59
69
74
79
83
Stimulus number

#### Figure S7

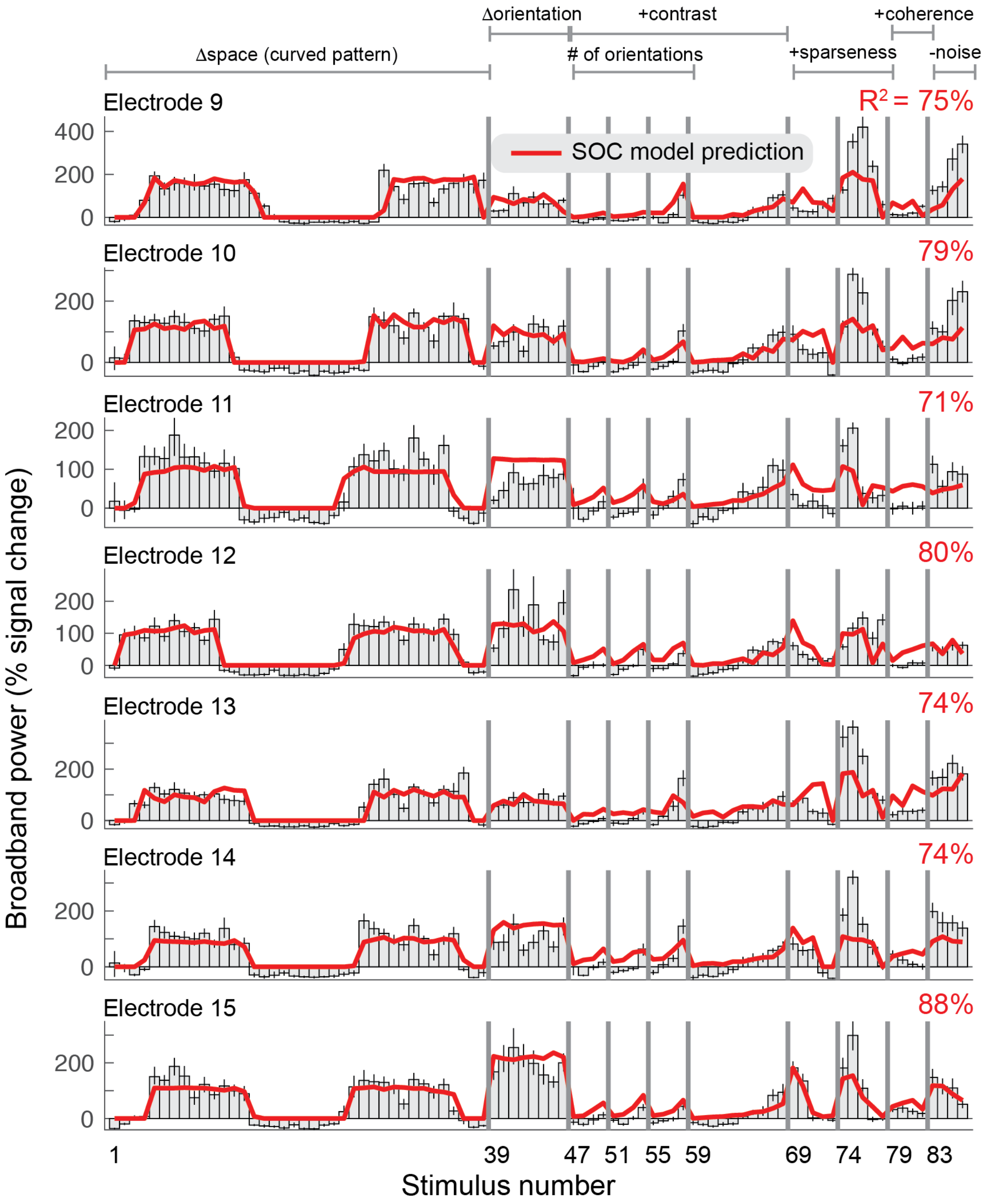


**Figure S6-S7. Second-order contrast (SOC) model accounts for ECoG broadband responses.** Each row shows the percent signal change in ECoG broadband power for all 86 stimuli for the 15 electrodes on V1, V2 or V3. Error bars display the 68% range for bootstrapped responses (bootstrapped across repeated presentations of the same stimuli). The SOC model was fit to these data using leave-one-stimulus-out cross-validation. The cross-validated predictions and amount of variance explained are shown in red.

#### Figure S8

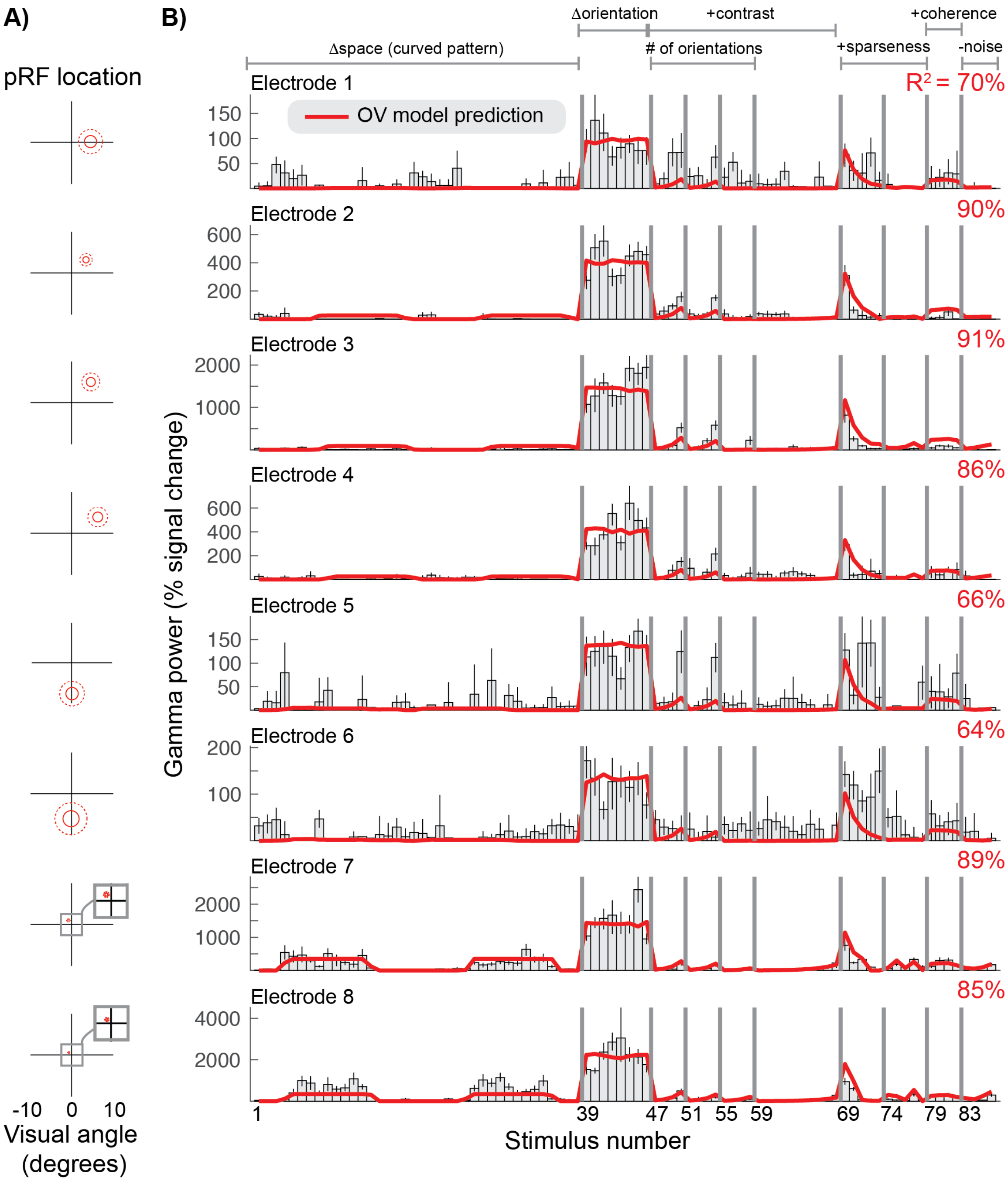
A)
B)
pRF location
-10 0 10
Visual angle (degrees)
Δorientation
+contrast
+coherence
Δspace (curved pattern)
### of orientations
+sparseness
-noise
Electrode 1
OV model prediction
$R^2$ = 70%
Electrode 2
90%
Electrode 3
91%
Electrode 4
86%
Electrode 5
66%
Electrode 6
64%
Electrode 7
89%
Electrode 8
85%
Gamma power (% signal change)
1 39 47 51 55 59 69 74 79 83
Stimulus number

#### Figure S9

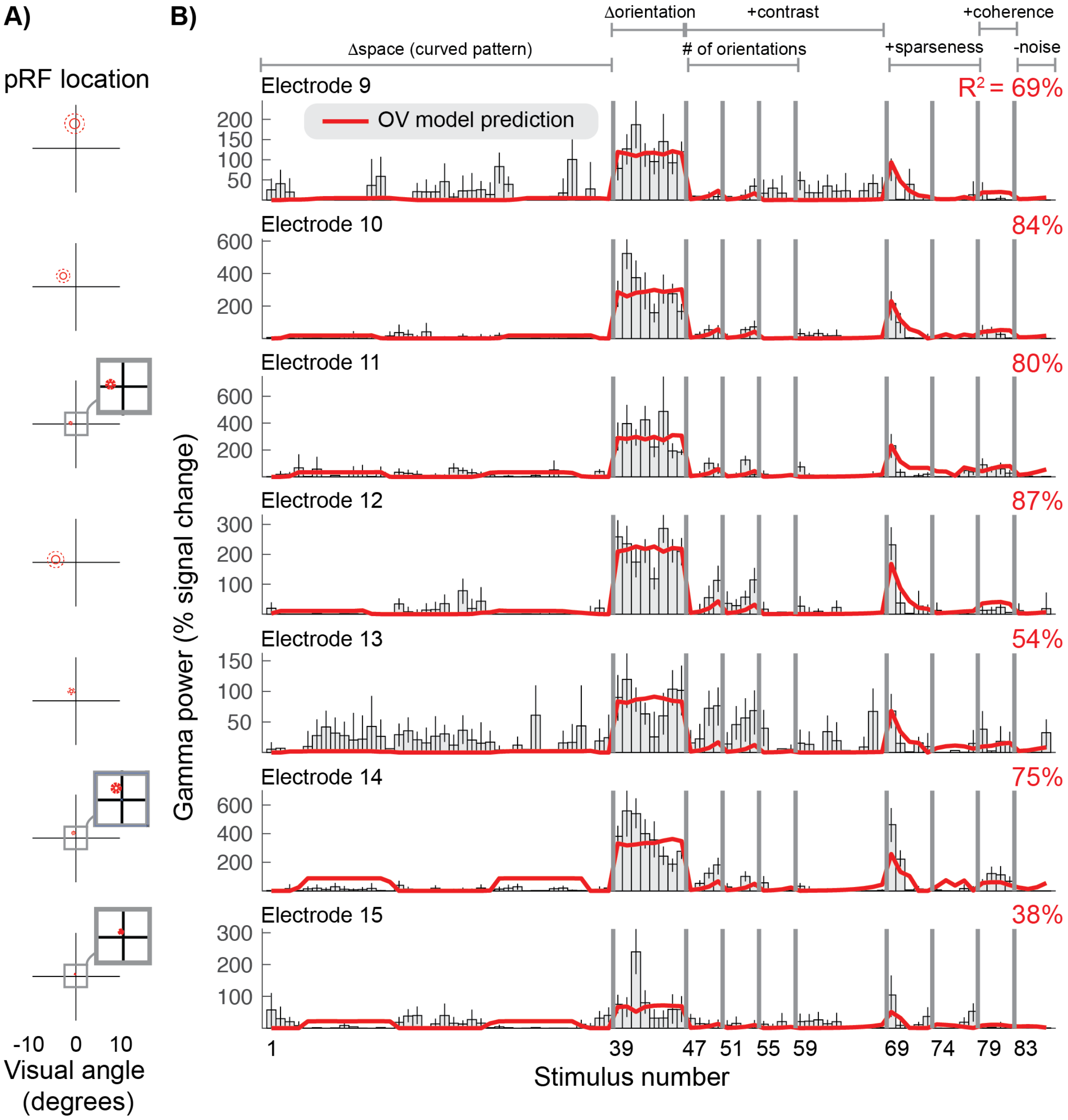


**Figure S8-S9. Orientation Variance (OV) model predicts selectivity of gamma responses. A)** The population receptive field for each electrode was defined by a Gaussian, indicated by the 1- and 2-sd contours (solid and dotted red lines). **B)** The gamma power in percent signal change was calculated for the 15 electrodes (rows) for all 86 stimuli. Error bars display the 68% range for the bootstrapped responses. The cross-validated predictions of the OV model and overall variance explained ($R^2$) are shown in red.

#### Figure S10

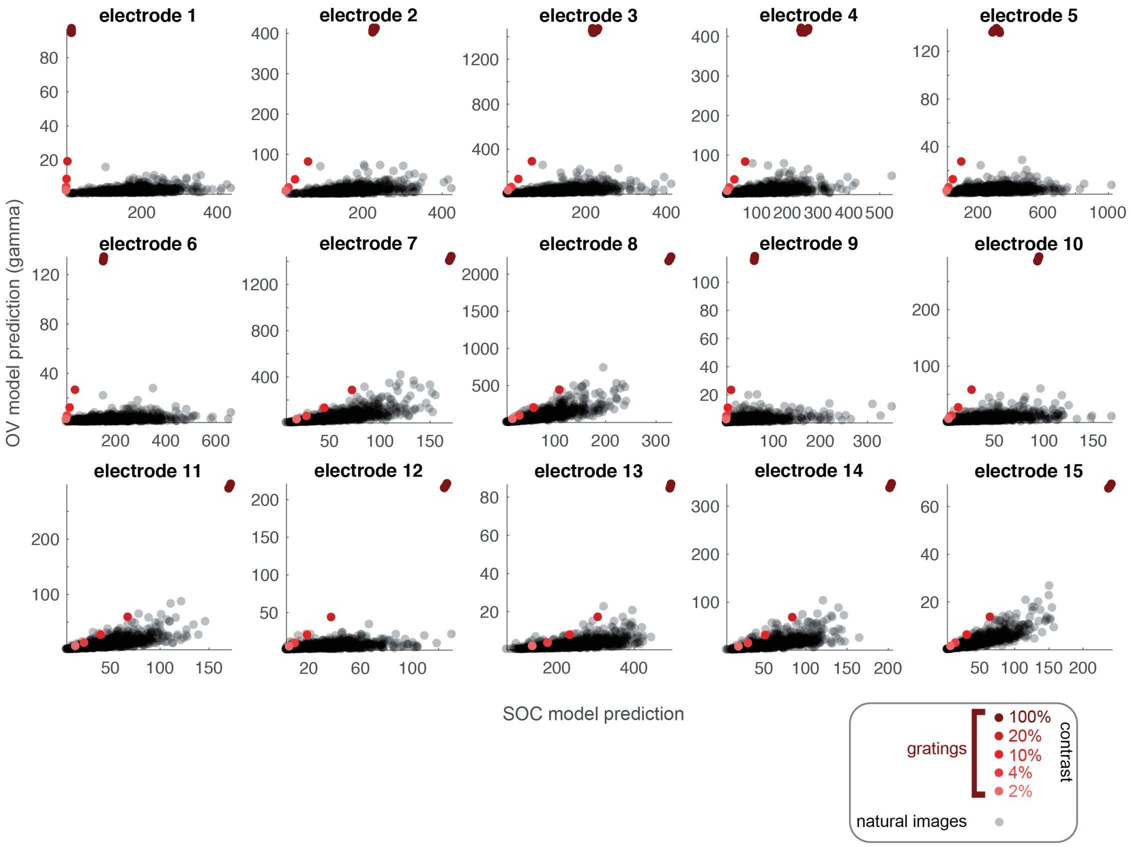


**Figure S10. OV and SOC model predictions for images of natural scenes for all electrodes.** The OV and SOC outputs are calculated for a set of gray-scale photographs of scenes, with model parameters from each electrode. The units are in percent signal change. Each gray dot is the output of the two models for one image. The red dots are the model outputs for grating stimuli of varying contrast. The cluster of red dots at 100% contrast displays high-contrast gratings of different orientations (stimuli 39-46).
